# Arginine-GlcNAcylation of death domain and NleB/SseK proteins is crucial for bacteria pathogenesis by regulating host cell death

**DOI:** 10.1101/746883

**Authors:** Juan Xue, Xing Pan, Lijie Du, Xiaohui Zhuang, Xiaobin Cai, Shan Li

## Abstract

Death receptor signaling is critical for cell death, inflammation, and immune homeostasis. Hijacking death receptors and their corresponding adaptors through type III secretion system (T3SS) effectors has been evolved to be a bacterial evasion strategy. NleB from enteropathogenic *Escherichia coli* (EPEC) and SseK1/2/3 from *Salmonella enterica* serovar Typhimurium (*S.* Typhimurium) can modify some death domains involved in death receptor signaling through arginine-GlcNAcylation. This study applied a limited substrate screen from 12 death domain proteins with conserved arginines during EPEC and *Salmonella* infection and found that NleB from EPEC hijacked death receptor signaling tumor necrosis factor receptor 1 (TNFR1)-associated death domain protein (TRADD), FAS-associated death domain protein (FADD), and receptor-interacting serine/threonine-protein kinase 1 (RIPK1), whereas SseK1 and SseK3 disturbed TNFR signaling through the modification of TRADD Arg235/245 and TNFR1 Arg376, respectively. SseK1 and SseK3 delivered by *Salmonella* inhibited TNF-α- but not TNF-related apoptosis-inducing ligand (TRAIL)-induced cell death, which was consistent with their host substrate recognition specificity. Taking advantage of the substrate specificity of SseK effectors, we found that only SseK1 fully rescued the bacteria colonization deficiency contributed by NleBc in *Citrobacter rodentium* infection animal model, indicating that TRADD was likely to be the preferred *in vivo* substrate corresponding to NleB/SseK1-induced bacterial virulence. Furthermore, novel auto-arginine-GlcNAcylation was observed in NleB and SseK1/3, which promoted the enzyme activity. These findings suggest that arginine-GlcNAcylation in death domains and auto-arginine-GlcNAcylation catalyzed by type III-translocated bacterial effector proteins NleB/SseKs are crucial for bacteria pathogenesis in regulating nuclear factor-κB (NF-κB) and death receptor signaling pathways. This study provides an insight into the mechanism by which EPEC and *Salmonella* manipulate death receptor signaling and evade host immune defense through T3SS effectors.

**Author Summary:** Enteropathogenic *Escherichia coli* (EPEC) and *Salmonella enterica* serovar Typhimurium (*S.* Typhimurium) are important food-borne pathogens infecting the intestine. They deliver type III secretion system effector NleB/SseKs to modify host death domain proteins by arginine GlcNAcylation. We screened the modification of 12 death domains containing conserved arginine in human genome by NleB, SseK1, SseK2, and SseK3 through ectopic co-expression and bacterial infection. Unlike multiple death receptor signaling inhibition by NleB, we found that SseK1 and SseK3 specifically hijacked tumor necrosis factor receptor 1 (TNFR1)-mediated death signaling through targeting TNFR1-associated death domain protein (TRADD) and receptor TNFR1, respectively. We identified the modification sites and suggested that TRADD was the *in vivo* target of NleB in mice infection model by utilizing the substrate specificity of SseK1 and SseK3, which highlighted anti-bacterial infection role of TRADD in death receptor signaling and non-death receptor signaling. In addition to the modification on host death domain substrates, we firstly elucidated the effect of auto-modification of the arginine GlcNAc transferases on the enzymatic activity, which widened our understanding of the newly discovered post translational modification in the process of pathogen-host interaction.

## Introduction

Death receptor signaling is crucial for cell death(1–3), inflammation(4), and immune homeostasis(5). Death receptor signaling is meditated by homotypic or heterotypic interactions among death domains (DDs) of the tumor necrosis factor receptor (TNFR) family of transmembrane death receptors and the downstream adaptors including TNFR1-associated death domain protein (TRADD)(3, 6), receptor-interacting serine/threonine-protein kinase 1 (RIPK1)(3, 7), and FAS-associated death domain protein (FADD)(3, 8). TRADD is known as the initial adaptor for TNFR1-induced apoptosis and nuclear factor-κB (NF-κB) signaling(6, 9–13). Recent studies have found that TRADD has Goldilocks effect on the survival of *Ripk1^−/−^Ripk3^−/−^* mice, and that *Tradd^+/+^* and *Tradd^−/−^* result in death through apoptosis, while haplosufficiency of TRADD is optimal for survival(14). RIPK1 and TRADD are synergisitically required for TNF-related apoptosis-inducing ligand (TRAIL)-induced NF-κB signaling and TNFR1-induced NF-κB signaling and apoptosis(15). FADD mediates TRAIL-induced necroptosis but antagonizes TNF-induced necroptosis(15–19). Type III Secretion system effector NleB from enteropathogenic *E.coli* (EPEC) was previously reported as a novel GlcNAc transferase that inhibited multiple death receptor mediated inflammation and cell death by modifying a conserved arginine residue in some death domain proteins(20–22). The arginine GlcNAc transferase activity of NleB is critical for A/E pathogen colonization in the mouse colon(20–23). Although modification of TRADD, FADD, and RIPK1 in *in vitro* reconstitution system and *ex vivo* epithelial cell infection system has been studied, the *in vivo* substrate preference of NleB remains elusive.

Arg-GlcNAcylation, a previously unappreciated post-translational modification, is not unique to extracellular bacteria pathogen EPEC(20, 22). Intracellular pathogen *Salmonella entrica* strains secreted three *Salmonella* pathogenicity island 2 (SPI-2) effector SseK1, SseK2 and SseK3(24–27). Crystal structure studies show that NleB, SseK1, and SseK3 belong to the GT-A family glycosyltransferase(22, 28, 29). The crystal structure of NleB in complex with FADD-DD and the sugar donor, NleB-GlcNAcylated death domains (TRADD-DD and RIPK1-DD) show that NleB is an inverting enzyme that converts the α-configuration in the UDP-GlcNAc donor into the β-configuration towards the conserved arginine of death domain proteins, namely, TRADD Arg235, FADD Arg117, and RIPK1 Arg603(22). Previous *in vitro* studies have suggested that SseK1 could GlcNAcylate TRADD(20), FADD(26), and GAPDH(27) with different efficiency, and that SseK3 could bind but not modify TRIM32(30), while it could slightly modify TRADD(26). One recent proteomics study has indicated the preferential substrate(s) is TRADD for SseK1, and TNFR1 and TRAILR for SseK3 during *Salmonella* infection in macrophage cells, however the substrate detection in their study highly depends on the endogenous protein abundance and mass spec performance(31). So the study of substrate specificity screening of SseK1 and SseK3 among death domain proteins during *Salmonella* infection is lacking, and the preferred *in vivo* substrates of the SseK effectors remain controversial.

Therefore, this study applied a limited substrate screen of 12 conserved arginine-containing death domain proteins during EPEC and *Salmonella* infection, finding that NleB from EPEC hijacked death receptor signaling via TRADD, FADD, and RIPK1, whereas SseK1 and SseK3 disturbed TNFR signaling by modifying TRADD and TNFR1, respectively. SseK1 GlcNAcylated hTRADD at Arg235 and Arg245 while SseK3 targeted TNFR1 at Arg376. SseK1 inhibited TRADD-activated NF-κB and apoptosis. SseK1 and SseK3 delivered by *Salmonella* inhibited TNF-α- but not TRAIL-induced cell death, which was consistent with their host substrate recognition specificities. Through the substrate specificity of SseK effectors, we found that only SseK1 fully rescued the bacteria colonization deficiency contributed by NleBc in *C. rodentium* infection animal model, indicating that TRADD was the possible *in vivo* substrate corresponding to NleB/SseK1-induced bacterial virulence. Furthermore, novel auto-arginine-GlcNAcylation was observed in NleB and SseK1/3, which promoted the enzyme activity. All these findings suggest that arginine-GlcNAcylation in death domains and auto-arginine-GlcNAcylation catalyzed by type III-translocated bacterial effector proteins NleB/SseKs are crucial for bacteria pathogenesis in regulating NF-κB and death receptor signaling pathways, which provides a better understanding of how EPEC and *Salmonella* using T3SS effectors to manipulate death receptor signaling and evade host immune defense.

## Results

### SseK1 and SseK3 hijack death receptor signaling via TRADD and TNFR1 respectively during *Salmonella* infection

Death domain (DD) is a subclass of protein motifs known as the death fold, thus DD-containing proteins are usually associated with programmed cell death and inflammation(4, 32, 33). Multiple sequence alignment of DDs in human genome revealed that Arg235 in TRADD (Arg117 in FADD) was conserved in one-third of the total 37 DD-containing proteins, including TNFR1, TRADD, FADD, RIPK1, FAS, DR3, DR4, DR5, ANK1, ANK2, ANK3, and ANKDD1B (Fig 1A and 1B, S1A Fig). Phylogenetic tree of the 12 DDs showed evolutionary relationship based on protein sequence (Fig 1C). Previous study indicated that the conserved arginine was critical for NleB modification(20, 22). To identify the substrate(s) of SseK family proteins, we screened all the arginine-containing death domain proteins both in biochemical reaction and co-expression systems. Most of the tested death domain proteins were modified by SseK1 and SseK3, and SseK2 exhibited weak enzymatic activity even in overexpression system as previously reported (S2 Fig)(26). To further identify the physiological substrates of these arginine GlcNAc transferases in bacterial infection process, we generated arginine GlcNAc transferase-deficient strains from EPEC and *Salmonella* and thus expressing NleB, SseK1, SseK2, and SseK3 individually in respected strains. NleB from EPEC E2348/69 modified TRADD DD, FADD, and RIPK1 DD (Fig 1D, S1B Fig). SseK1 and SseK3 from *Salmonella* Typhimurine SL1344 specifically modified TRADD DD and TNFR1 DD, respectively, while SseK2 exhibited no obvious GlcNAcylation activity towards death domain proteins during infection (Fig 1E-1G, S1B and S3 Figs).

**Fig 1.**
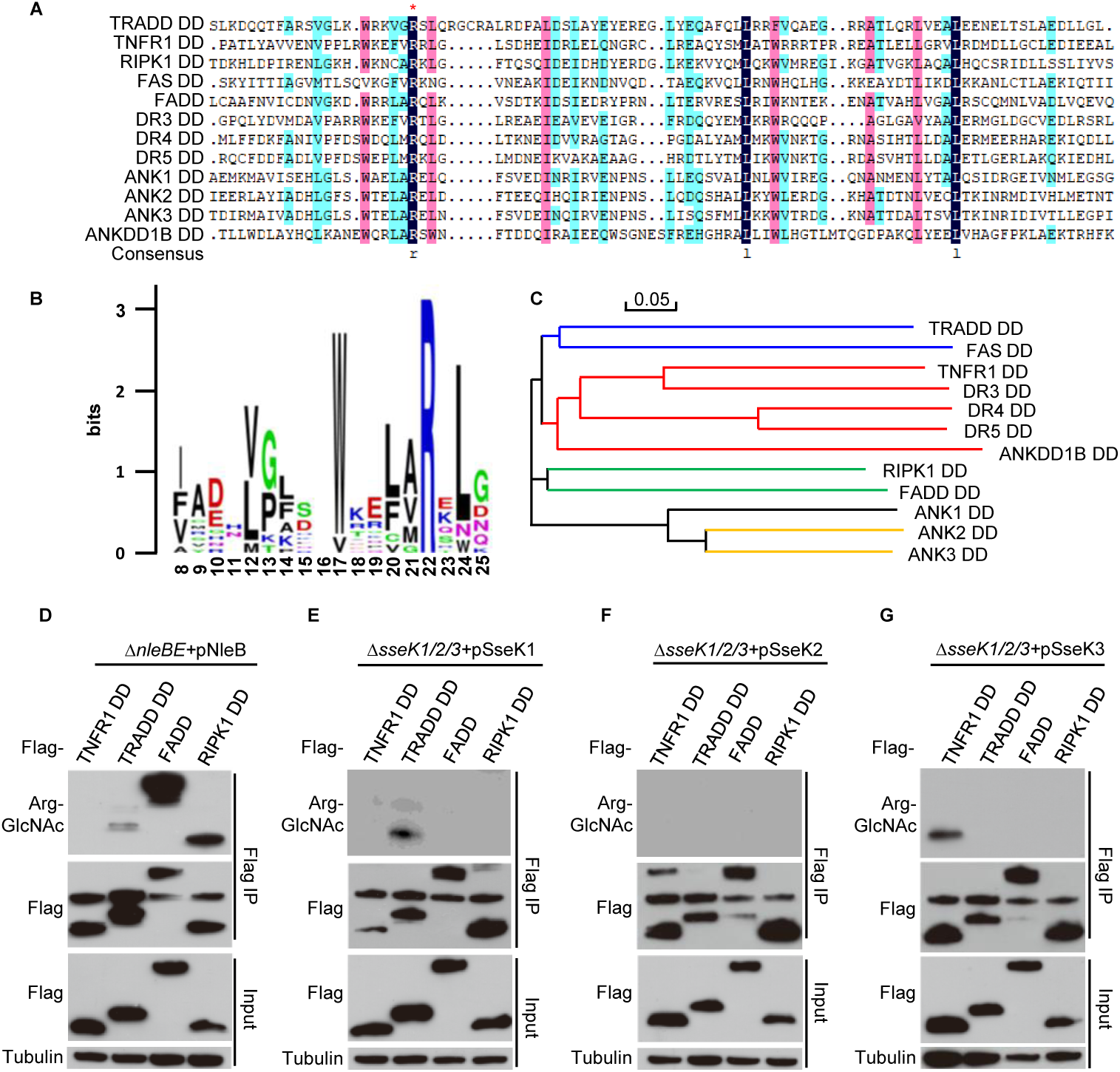
SseK1 and SseK3 hijack death receptor signaling via TRADD and TNFR1, respectively during *Salmonella* infection. (A) Multiple sequence alignment of 12 death domains (DDs) from human proteins containing death domain. The red asterisk indicates the conserved arginine site among different DDs, which could be GlcNAcylated by NleB/SseKs. **(B)** Site distribution of amino acids in 12 death receptor domains containing conserved arginine by Weblogo. **(C)** Phylogenetic tree of the 12 DDs constructed on the basis of sequence similarity. **(D-G)** Identification of the physiological substrates of arginine GlcNAc transferase NleB/SseKs during bacterial infection of mammalian cells. 293T cells transfected with pCS2-1Flag-TNFR1 DD, pCS2-1Flag-TRADD DD, pCS2-1Flag-FADD, and pCS2-1Flag-RIPK1 DD were infected with the indicated modified EPEC strains or *Salmonella* strains. After infection, cells were lysed, and proteins were immunoprecipitated with FLAG M2 beads. Samples were loaded onto SDS-PAGE gels and were immunoblotted with anti-Flag, anti-Arg-GlcNAc, and a loading control anti-tubulin. Data in Fig **(D**-**G)** are from at least three independent experiments.

**Fig 2.**
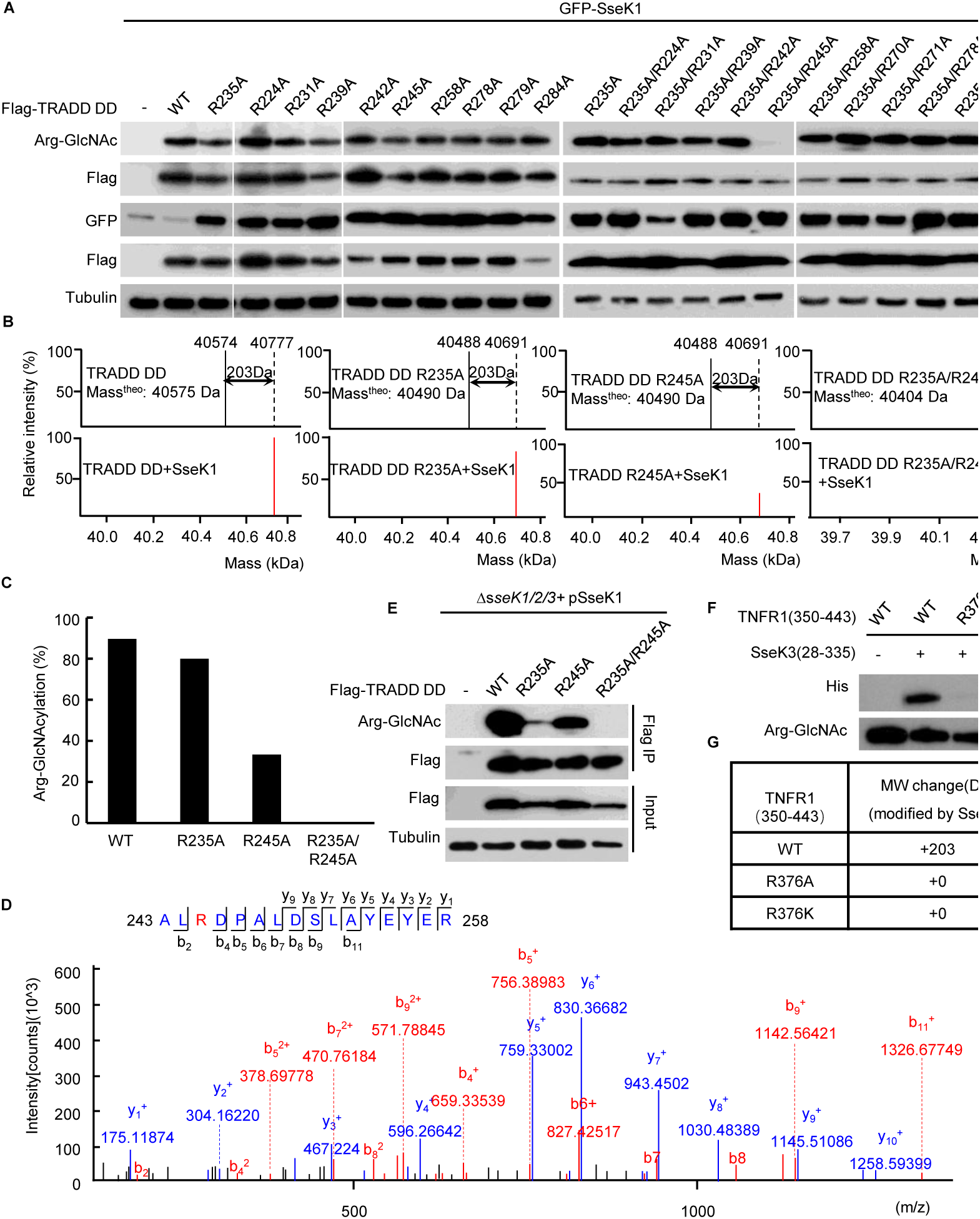
SseK1 GlcNAcylates hTRADD at R235/R245 and SseK3 GlcNAcylates TNFR1 at R376. (A) An arginine point mutation screen of hTRADD to investigate its ability to be GlcNAcylated by SseK1. 293T cells were transfected with the indicated plasmid combinations. The samples of anti-Flag immunoprecipitates (Flag IP) and total cell lysates (Input) were immunoblotted with corresponding antibodies. Anti-tubulin was used as a loading control. **(B)** Electrospray ionization mass spectrometry (ESI-MS) determination of the total mass of the site-directed TRADD mutants purified from bacteria. GST-TRADD DD, GST-TRADD DD (R235A), GST-TRADD DD (R245A), and GST-TRADD DD (R235A/R245A) were expressed alone (upper panel) or co-expressed with His-SseK1 (lower panel) in *E. coli* BL21 (DE3) strain. The resulting mass spectra were shown. The resulting mass spectra were shown. The black bar and red bar denote unmodified and GlcNAcylated TRADD DD, respectively. **(C)** The percentage of site-directed TRADD mutants GlcNAcylated by SseK1. **(D)** HCD analysis of the peptides of TRADD DD R245 GlcNAcylated by SseK1 in bacteria. The fragmentation patterns of the generated b and y ions were shown along the peptide sequence on the top of the spectrum. **(E)** Modification of TRADD and TRADD variants by SseK1 upon *S.* Typhimurine infection. 293T cells was transfected with plasmids carrying TRADD and the site-directed TRADD mutants, and then infected with the indicated *Salmonella* strains. After 15-hour infection, cells were lysed and proteins were immunoprecipitated with FLAG M2 beads. Samples were loaded onto SDS-PAGE gels, followed by immunoblots with anti-Flag and anti-Arg-GlcNAc antibodies. **(F)** Identification of the site of TNFR1 GlcNAcylated by SseK3 in bacteria. **(G)** Summary of ESI-MS determination of the total mass of TNFR1 and its point mutants co-expressed with SseK3 in bacteria. Data in Fig **(A, E,** and **F)** are from at least three independent experiments.

**Fig 3.**
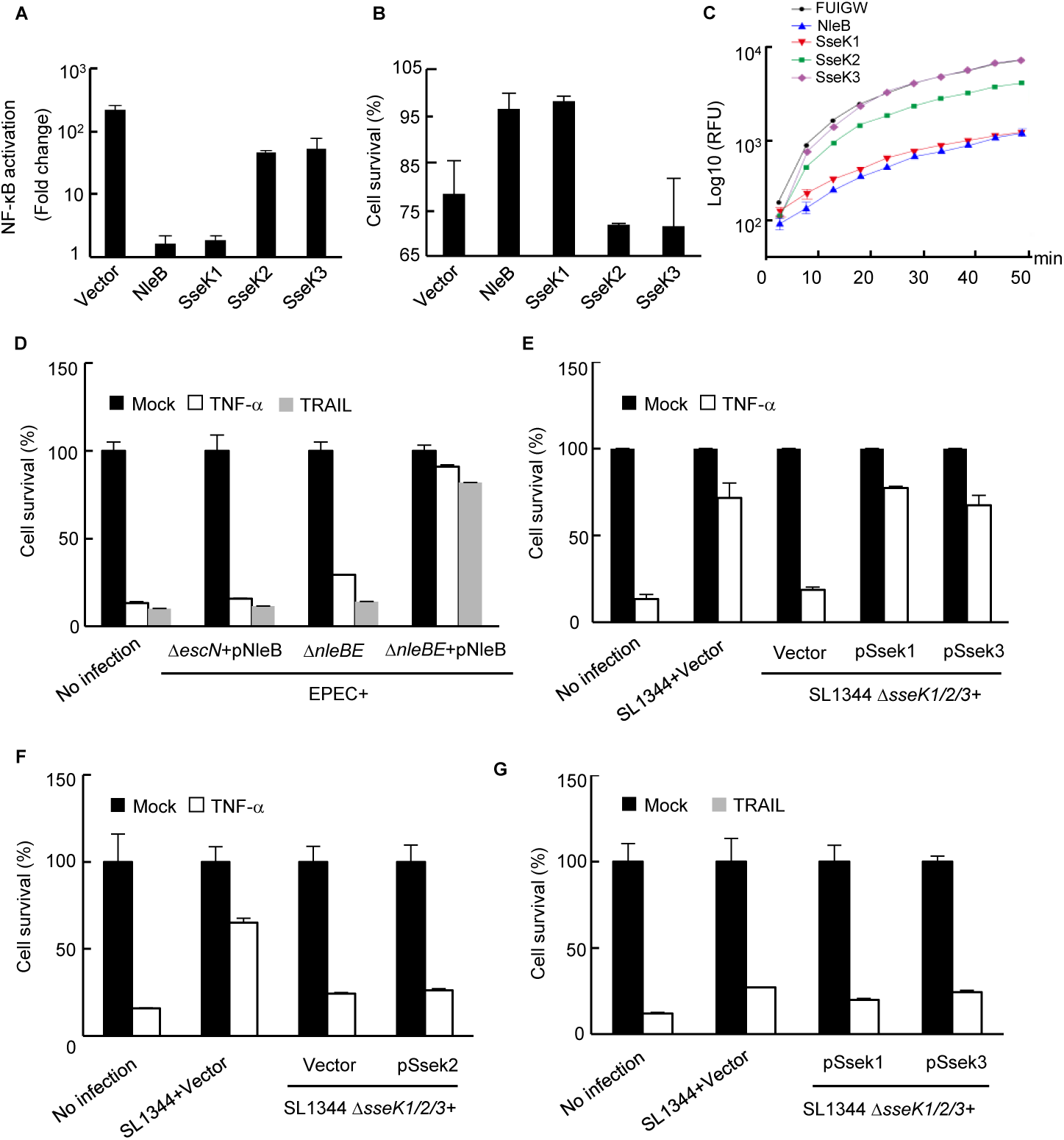
SseK1 rather than SseK3 inhibits TRADD-activated NF-κB and cell death signaling in 293T cells, and both SseK1 and SseK3 inhibit TNF-α-induced but not TRAIL-induced cell death during *Salmonella* infection. **(A-C)** Effects of NleB/Sseks on TRADD DD-activated NF-κB **(A)**, cell death **(B)**, and caspase-3 activation **(C)**. 293T cells were transfected with Flag-TRADD DD in combination with the indicated plasmids. NF-κB luciferase activation was indicated as fold change **(A)**. Cell viability was determined by measuring ATP levels **(B)**. Caspase-3 activity was assayed using an Ac-DEVD-AFC probe according to manufacturer’s instructions **(C)**. **(D-G)** HeLa cells infected with the indicated EPEC strains and *Salmonella* strains were stimulated with TNF-α and TRAIL. Cell viability was determined by measuring ATP levels. Data were normalized according to the uninfected cell data. Black bars or white bars denoted the unstimulated or stimulated cells, respectively. Data in (**A-G)** were derived from at least three independent experiments.

### SseK1 GlcNAcylates hTRADD at Arg235/Arg245 and SseK3 GlcNAcylates TNFR1 at Arg376

We found that the mutation of Arg235, which was the only modification site on TRADD by NleB(20), could not abolish the modification of human TRADD by SseK1, which was in accordance with previous report on mouse TRADD(26), suggesting that SseK1-mediated GlcNAcylation of TRADD occurred on either a different amino acid or on multiple arginine residues. To fully explore the modification site(s) of TRADD by SseK1, we applied an arginine point mutation screen of hTRADD to investigate its ability to be GlcNAcylated by SseK1. We transfected the HEK293T cells with pEFGP-SseK1 individually or together with wild type Flag-hTRADD or with different point mutation variants. Single point mutation of any arginine in the death domain of TRADD did not abolish the modification (Fig 2A), which indicated the existence of multiple modification sites. Subsequently, we mutated all the arginine residues against the background of Arg235Ala mutation. By contrast to wild type TRADD, the GlcNAcylation of hTRADD at Arg235Ala/Arg245Ala was significantly reduced upon arginine-GlcNAcylation specific antibody detection (Fig 2A). To determine the modification ratio, recombinant GST-TRADD DD and its point mutations were expressed alone (upper panel in Fig 2B) or co-expressed with SseK1 (lower panel in Fig 2B) in *E. coli* BL21 (DE3) strain. Purified proteins were analyzed on the mass spectrometer and the resulting total molecular weight of mass spectra was shown in Fig 2B. Electrospray ionizations mass spectrometry (ESI-MS) analysis identified a 203-dalton (Da) increase in the total mass of TRADD DD, TRADD DD at Arg235Ala, and TRADD DD at Arg245Ala co-expressed with SseK1, whereas TRADD DD at Arg235Ala/Arg245Ala co-expressed with SseK1 merely showed the theoretic molecular mass, which was consistent with other TRADD DD proteins expressed alone (Figs 2B and 2C). LC-MS/MS analysis revealed that the GlcNAc modification occurred at Arg224, Arg242, and Arg245 (Fig 2D, S4 Fig), as well as Arg235(26). Considering that the TRADD DD (Arg235Ala/Arg245Ala) mutant exhibited no 203 Da molecular weight increase upon SseK1 treatment, Arg235 and Arg245 might be the dominant modification sites. Consistently, SseK1 delivered from a *Salmonella* derivative strain SL1344Δ*sseK1/2/3* by type III secretion system, could not modify TRADD DD at R235A/R245A yet, even though single arginine mutation could be modified (Fig 2E). For the modification of TNFR1 by SseK3, mutation of Arg376 into Ala or Lys abolished the arginine-GlcNAcylation signal and molecular weight increase, suggesting that the conserved Arg376 was the *bona fide* modification site (Fig 2F and 2G). All these data showed that SseK1 GlcNAcylated hTRADD at R235 and R245 both in co-expression system and in pathogen infection process. SseK3 modified TNFR1 at the predicted R376 residue.

**Fig 4.**
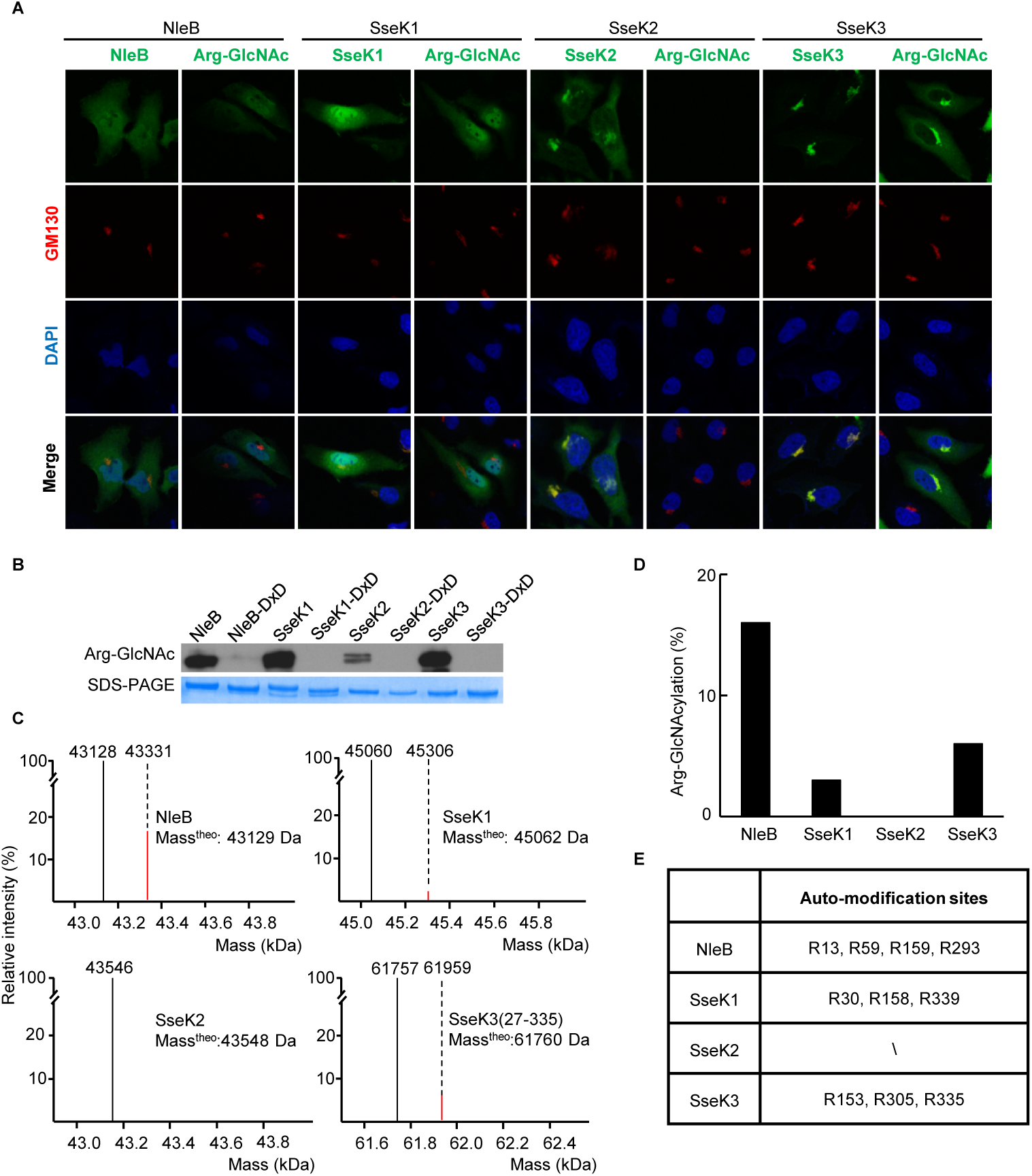
Novel auto-arginine-GlcNAcylation observed in NleB and SseKs. (A) Ectopic expression of NleB/SseKs effectors showed the related subcellular localization and modification pattern in transfected HeLa cells. GFP-NleB, GFP-SseK1, GFP-SseK2, and GFP-SseK3 were expressed ectopically in HeLa cells. In HeLa cells, green indicated immunofluorescence staining of GFP and arginine-GlcNAcylated proteins. Blue indicated DAPI staining of nuclei, and red indicated GM130 staining of the Golgi structure. **(B)** Analysis of NleB/SseKs auto-arginine-GlcNAcylation by Western blot. Recombinant purified NleB/SseKs and their enzymatic mutants were analyzed on SDS-PAGE gels, followed by immunoblotting with anti-Arg-GlcNAc. **(C)** ESI-MS analysis determination of the total mass of the NleB/SseKs purified from bacteria. The black bar denotes unmodified protein. For NleB and SseK3, the red bar denotes GlcNAcylated form with 203-Da increase, while for SseK1, the red bar denotes GlcNAcylated (203 Da) and acetylated (42 Da) form with 245-Da increase. **(D)** Arginine-GlcNAcylation percentage of NleB/SseKs. **(E)** Summary of ESI-MS determination of the total mass of NleB/SseKs. Data in **(A** and **B)** are representative from at least three independent experiments.

### SseK1 rather than SseK3 inhibits TRADD-activated NF-κB and cell death signaling in 293T cells

It has been reported that robust inhibition of TNF-α-stimulated NF-κB signaling can be blocked by ectopic expression of SseK1 and SseK3 in various cell lines(26). Similar to the observation in TNF-α stimulation, SseK1 completely abolished TRADD overexpression-induced NF-κB activation (Fig 3A) and apoptosis in 293T cells (Fig 3B and 3C). Thus, SseK1 could target TRADD and disrupt multiple signaling pathways at the downstream of TNFR1.

### SseK1 and SseK3 inhibit TNF-α-induced but not TRAIL-induced cell death during *Salmonella* infection

The death domains of TRADD, FADD, and RIPK1 could be GlcNAcylated by NleB, leading to the inhibition of multiple death receptor-mediated cell death as well as NF-κB signaling(20, 22). Considering that SseK1 and SseK3 could only modify TRADD DD and TNFR1 DD during infection, respectively, we tested whether SseK proteins could specifically inhibit certain death receptor-induced cell death. First, we confirmed that NleB delivered by EPEC almost completely inhibited both TNF-α-induced and TRAIL-induced cell death (Fig 3D). Next, we analyzed the cell survival percentage of HeLa cells firstly infected with the indicated *Salmonella* derivative mutant strains, subsequently treated with TNF-α or TRAIL or without treatment. Cell viability was determined by measuring ATP levels. We found that the infection with *Salmonella* strain SL1344 wild type strain inhibited TNF-α-induced but not TRAIL-induced cell death. Simultaneous deletion of SseK1, SseK2, and SseK3 abolished the inhibitory effect (Fig 3E-3G). Rescue of SseK1 or SseK3 but not SseK2 in the triple deletion mutant Δ*sseK1/2/3* was sufficient to inhibit TNF-α but not TRAIL-induced cell death (Fig 3E-3G), which was consistent with the substrate specificity under infection condition.

### Novel auto-arginine-GlcNAcylation is observed in NleB and SseKs

When we detected the subcellular localization of NleB/SseKs and their arginine GlcNAcylation in HeLa cells, we found the related subcellular localization and modification pattern through the ectopic expression of NleB/SseKs effectors. GFP-NleB and GFP-SseK1 were diffusely localized in cytoplasm, and the respected arginine GlcNAcylation showed no specific subcellular localization in the transfected HeLa cells (Fig 4A). The arginine GlcNAcylation was not detected in SseK2-transfected cells due to the weak GlcNAc transferase activity, whereas the GFP-SseK2 was found to be co-localized with cis-Golgi marker GM130 (Fig 4A). GFP-SseK3 and its arginine GlcNAcylation were co-localized in the host Golgi network (Fig 4A). Then, we detected the modification of NleB and SseKs. We found that these effectors could GlcNAcylate themselves when they were expressed as recombinant proteins both in prokaryotic and eukaryotic systems (Fig 4B and 5C). Mass spectrometry analysis determined the percentage of 203-Da increase in the total molecular weight of NleB, SseK1 and SseK3 (Fig 4C and 4D). SseK2 exhibited little mass change, which was in accordance with the weak signal detected by arginine GlcNAcylation antibody (Fig 4C and 4D). Arg13/59/159/293 in NleB, Arg30/158/339 in SseK1, and Arg153/305/335 in SseK3 are detected modification sites (Fig 4E, S5-S7 Figs), as well as Arg184(31). Interestingly, we also found that arginine GlcNAcylation widely existed in various pathogenic bacteria (S8 Fig).

**Fig 5.**
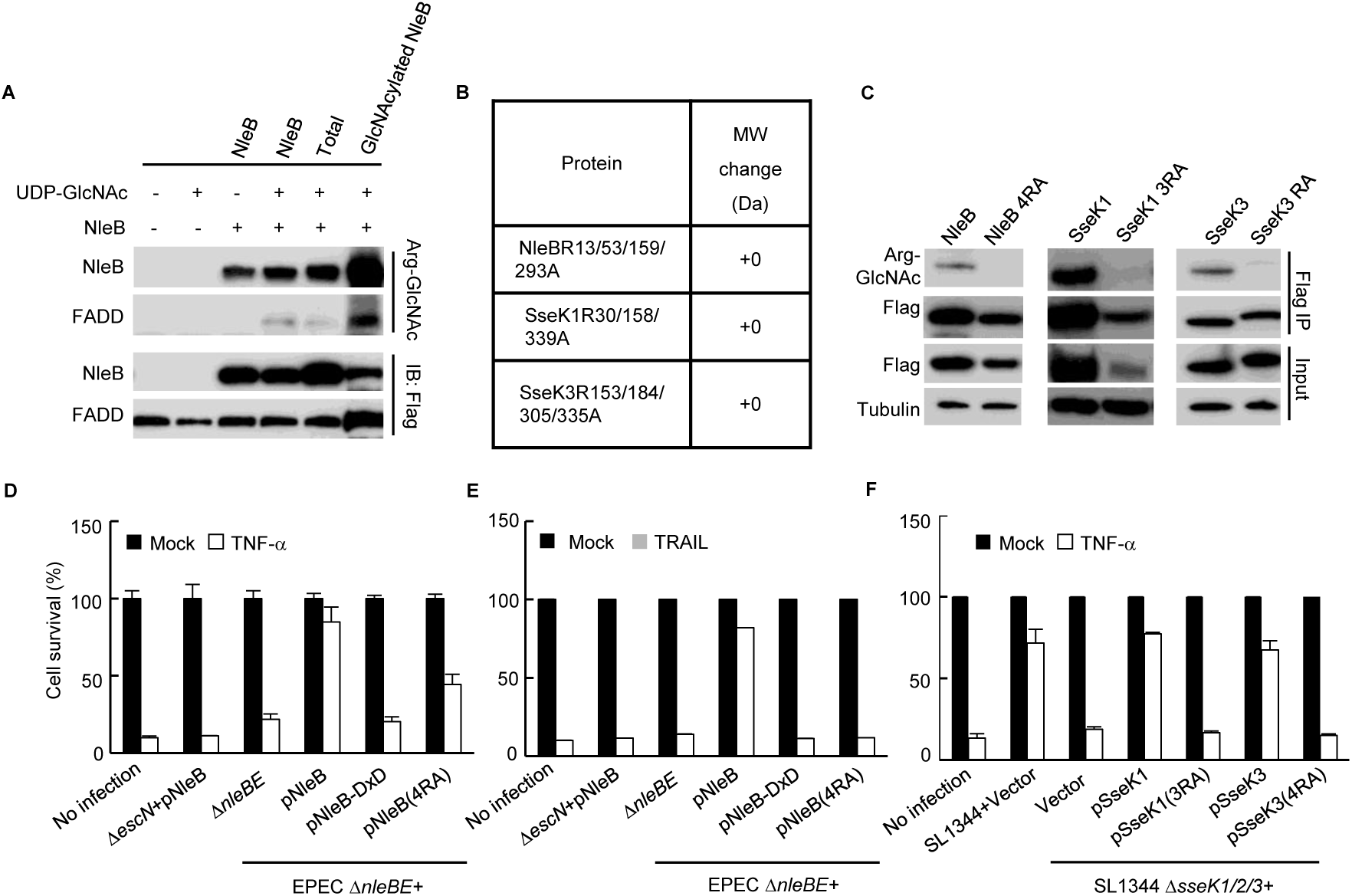
Auto-arginine-GlcNAcylation promotes the enzymatic activity of NleB and SseK1/3. (A) Effects of the auto-arginine-GlcNAcylation of NleB on enzyme activity towards death domain protein. The coupled anti-Arg-GlcNAc beads were incubated with 50 μg of purified NleB for enrichment with auto-arginine-glycosylated NleB. Beads enriched with auto-arginine-glycosylated proteins were used *in vitro* glycosylation assay. Samples were loaded onto SDS-PAGE gels and were immunoblotted with anti-Flag and anti-Arg-GlcNAc. **(B)** Mass spectrometry analysis of 203-Da increase in the total molecular weight of the auto-arginine-GlcNAcylation site-directed mutant proteins. **(C)** Effects of the modification site mutation of NleB/SseKs. 293T cells were transfected with the indicated plasmids. After 24-hour transfection, cells were lysed, and proteins were immunoprecipitated with α-FLAG conjugated beads. Samples were loaded onto SDS-PAGE gels, followed by immunoblot with anti-Flag, anti-Arg-GlcNAc, and a loading control anti-tubulin. **(D-F)** Effects of the auto-arginine-GlcNAcylation on cell death inhibition of NleB and SseKs. HeLa cells infected with the indicated EPEC strains and *Salmonella* strains were stimulated with TNF-α and TRAIL. Cell viability was determined by measuring ATP levels. Black bars or white bars denoted unstimulated or stimulated, respectively. Data in **(A)** and **(C-F)** are representative from at least three independent experiments.

### Auto-arginine-GlcNAcylation promotes the enzymatic activity of NleB and SseK1/3

To test whether auto-arginine-GlcNAcylation was functional, we enriched the GlcNAcylated NleB with the modification specific antibody and found that the auto-arginine-GlcNAcylation promoted the enzymatic activity of NleB towards death domain protein (Fig 5A). To investigate the effect of loss of auto-arginine-GlcNAcylation on NleB/SseK1/SseK3 transferase activity, site-directed recombinant proteins were expressed. ESI-MS analysis indicated that no 203 Da increase was observed, and that the mutants completely lost its auto-arginine-GlcNAcylation activity (Fig 5B and 5C). To validate its biological function, we tested whether loss of auto-arginine-GlcNAcylation of NleB/SseK1/SseK3 still inhibited TNF-α- or TRAIL-induced cell death during infection of HeLa cells. The survival cells were significantly reduced in HeLa cells infected with the mutants with auto-arginine-GlcNAcylation loss, compared to those in HeLa cells infected with wild type bacteria (Fig 5D-5F). Therefore, loss of auto-arginine-GlcNAcylation decreased the biological activity of NleB and SseK1/3.

### Only SseK1 fully rescues the bacteria colonization deficiency contributed by NleBc in *C. rodentium* infection animal model

Previous studies showed that arginine GlcNAc transferase activity of NleB was crucial for bacterial colonization and virulence in mice infection model(20–22). Considering the broad substrate range of NleB during *ex vivo* infection, it is intriguing to determine the major target(s) among TRADD, FADD, and RIPK1 in *in vivo* infection system. Complementation strains of Δ*nleB* with *Citrobacter* NleBc or *Salmonella* SseK1/2/3 replaced with signal peptide of NleBc were generated. MEF cells were firstly infected with the indicated *C. rodentium* strains, and then treated with CHX plus TNF-α. Subsequently, Cell viability was detected. The results indicated that complementation of Ssek1 and SseK3, but not SseK2, or respected DxD mutants, inhibited TNF-α-induced cell death, compared to the level of complementation of NleB in infected MEF cells, suggesting that chimera SseK1 and SseK3 were secreted and functioned in *Citrobacter rodentium* (Fig 6A). Afterwards, C57BL/6 mice were gavaged with *C. rodentium* derivatives. Complementation of Δ*nleB* with SseK1 rather than SseK2 or SseK3, recovered bacteria level in stool counts to the level of the strain complemented with NleB, and restored bacterial colonization in the intestinal tract of mice (Fig 6B). SseK1 exhibited substrate specificity to TRADD DD, which indicated TRADD DD was the preferential *in vivo* substrate corresponding to NleB-induced bacterial virulence (Fig 6B).

**Fig 6.**
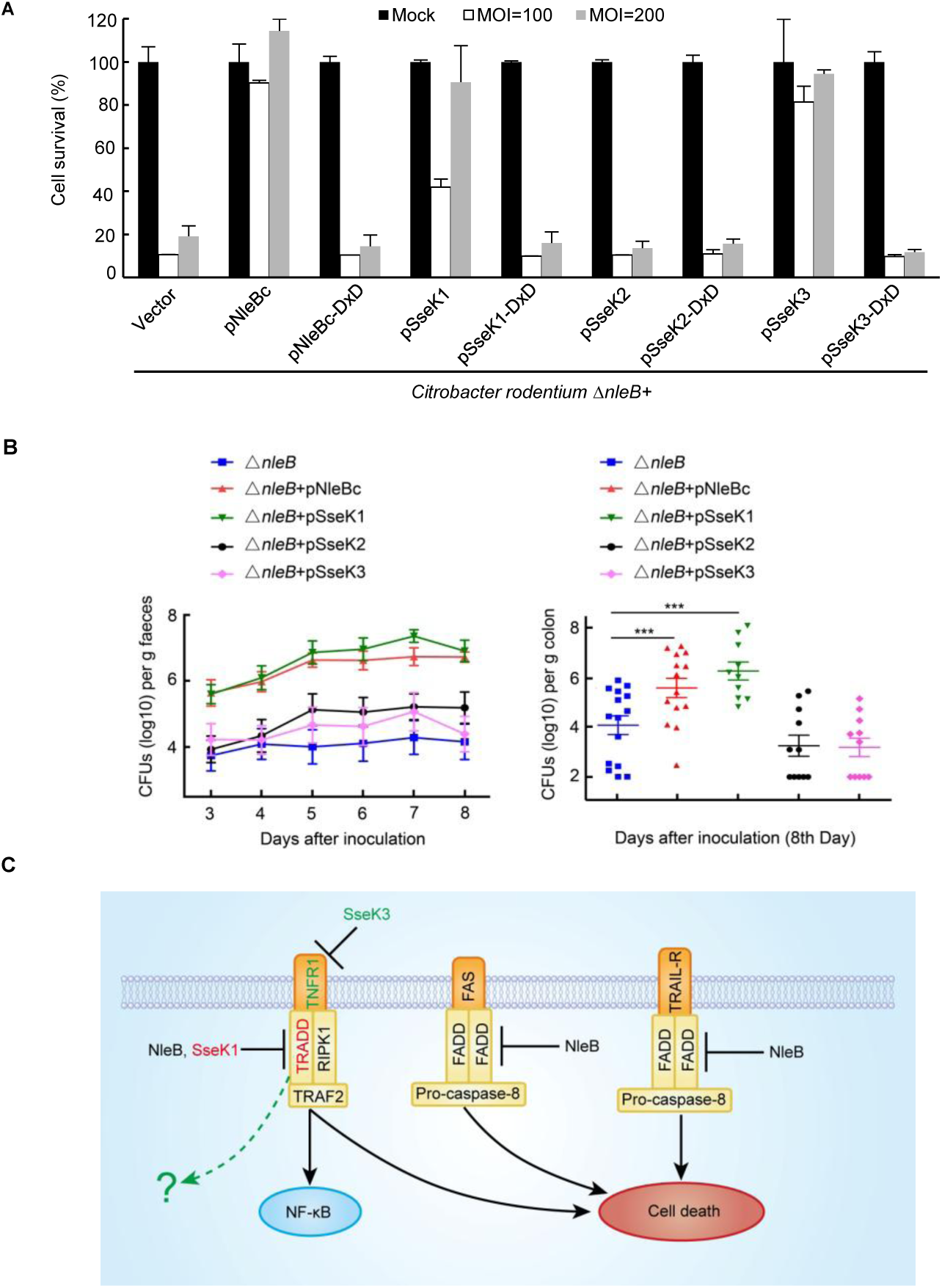
Only SseK1 fully rescues the bacteria colonization deficiency contributed by NleBc in *C. rodentium* infection animal model. **(A)** NleB and SseK1/3 inhibited TNF-α-induced cell death in MEF cells during *C. rodentium* infection. MEF cells infected with derivatives of *C. rodentium* strains were stimulated with TNF-α. Cell viability was determined by measuring ATP levels. Blue bars or red bars denoted unstimulated or stimulated, respectively. **(B)** Comparison of the colonization of different derivatives of the *C. rodentium* strains in mice colon after infection. 5-6-week-old C57BL/6 male mice were orally gavaged with *C. rodentium* derivatives. The viable stool bacteria count (log10 CFU/g feces) (left) and bacterial colonization in the intestine (log10 CFU/g colon, n >6) 8 days post infection (right) were presented as mean± s.e.m. \*\*\**P* < 0.001 was considered as the highest degree of significant difference. **(C)** Inhibition of death receptor (TNFR1, FAS and TRAIL-R) signaling by bacterial pathogen T3SS effectors NleB and SseKs. NleB was derived from enteropathogenic*. E coli* (EPEC) and enterohemorrhagic *E. coli* (EHEC). SseKs were homologues of NleB in *Salmonella*.

## Discussion

Previous studies confirmed that the arginine GlcNAc transferase activity of NleB was essential for bacterial colonization in the mouse model of EPEC infection(20–22). Considering that the death domains of TRADD, FADD, and RIPK1 could be modified by NleB in cell infection system(20, 22, 23, 34), it is necessary to determine the major target(s) *in vivo*. In this study, we found that only SseK1 fully rescued the bacteria colonization deficiency contributed by NleB from *Citrobacter* or EPEC in *C. rodentium* infection animal model. The finding that SseK1 had substrate specificity to TRADD indicated that TRADD was the preferential *in vivo* substrate corresponding to NleB-induced bacterial colonization, which might be correlated with the role of dynamic equilibrium of TRADD in both cell death and NF-κB signaling, and provide insights into the non-TNFR1 signaling role of TRADD (Fig 6C)(12, 13).

In addition to the Arg-GlcNAcylation of host death domain-containing proteins, we also observed the auto-arginine-GlcNAcylation of NleB, SseK1, and SseK3 by measuring total molecular weight by mass spec. Auto-modification of enzymes catalyzing post translational modifications usually played a significant role in self-activation, and it was indispensable in signaling transductions, such as autophosphorylation(35–38) and autoacetylation(39–45). Although the auto-Ser/Thr-GlcNAcylation on O-GlcNAc transferase (OGT)(46–48) and auto-arginine-GlcNAcylation on NleB(29) have been detected with antibodies(49–53) in western blot, the percentage and the physiological significance of the self-modification remain unknown. Here, we identified several auto-modification sites on NleB, SseK1, as well as Ssek3, finding that the auto-arginine-GlcNAcylation promoted the enzymatic activity of NleB towards FADD, which was crucial for bacterial infection.

Interestingly, using the specific Arg-GlcNAc antibody developed by our group, we also found that arginine GlcNAcylation widely existed in various pathogenic bacteria including *Escherichia coli, Shigella, Salmonella, Klebsiella, Burkholderia, Staphylococcus, Legionella, Citrobacter,* and *Vibrio*. Some of these bacteria may have NleB/SseK orthologs, some may not, suggesting that the modification is not unique. Therefore, a stimulating question arises whether arginine GlcNAcylation is generally involved in regulating cellular processes in prokaryotes and/or eukaryotes. Likewise, some researchers also observed arginine-GlcNAcylation on EPEC proteins and detected several arginine-GlcNAcylated proteins in macrophages during *Salmonella* infection(26).

In summary, this study presented compelling evidence for distinct substrate specificities of NleB, SseK1, SseK2, and SseK3, and further demonstrated that SseK1 and SseK3 specifically inhibited TNF-α but not TRAIL-induced cell death during *Salmonella* infection, and that only SseK1 fully rescued the bacteria colonization deficiency contributed by NleBc in *C.rodentium* infection animal model. Additionally, the existence of auto-arginine-GlcNAcylation of NleB and SseK1/3 was observed, which promoted the enzymatic activity of these T3SS effectors.

## Materials and Methods

### Bacteria strains and growth conditions

The EPEC strains, *Salmonella* strains, and *C. rodentium* strains used in this study are listed in S1-S3 Tables. The bacterial strains, unless specially mentioned, were grown in LB broth at 37°C with shaking in the presence of the following appropriate concentrations: nalidixic acid (50 μg/ml), kanamycin (50 μg/ml), ampicillin (100 μg/ml), chloramphenicol (17 μg/ml), and streptomycin (50 μg/ml).

### Plasmid construction and primers

All the plasmids and primers used in this study are listed in S4 and S5 Tables. DNA of NleB genes and NleB homologue genes was amplified from EPEC E2348/69, *C. rodentium* ICC168, and *S. enterica* Typhimurium SL1344 strains, and was inserted into pCS2-EGFP, pCS2-1Flag, and pCS2-3Flag for mammalian cell expression, and into pGEX-6P-2, pET28a-LFn, and pET28a-His for protein expression in *E. coli*. NleB and NleB homologue genes were ligated into the pTRC99A vector for complementation in EPEC (under the trc promoter) and into the pET28a vector for complementation in *C. rodentium* (under the *C. rodentium nleB* promoter). Human cDNAs TRADD, TNFR1, FADD, DR3DD, ANK1DD, ANKDD1B, ANK2DD, ANK3DD, RIPDD, FASDD, DR4DD, DR5DD were amplified from a HeLa cDNA library as previously described(20). DNA polymerases KOD Plus Neo (Toyobo, 536900) and Taq MasterMix (CWBIO, CW0682B) were used in accordance with the manufacturer’s instructions. All single point mutants were generated by quick change and multiple point mutants and truncation mutants were generated by standard molecular biology procedures. NF-κB reporter plasmids were used as previously described(20, 54). Plasmids were prepared by GoldHi endofree plasmid maxi kit (CWBIO, CW2104) or TIANGEN Spin Miniprep Kit. All plasmids were verified by DNA sequencing and primers were synthesized by Sangon Biotech.

### Antibodies and reagents

The anti-GlcNAc arginine antibody (ab195033, Abcam) was described previously(53). Antibody for Flag M2 (F2426) and tubulin (T5186) were Sigma products. Anti-EGFP (sc8334) was purchased from Santa Cruz Biotechnology, and anti-DnaK 8E2/2 (ab69617) was from Abcam. Horse radish peroxidase (HRP)-conjugated goat anti-mouse IgG (NA931V) and HRP-conjugated goat anti-rabbit IgG (NA934) were all purchased from GE Healthcare. Cell culture products were purchased from Invitrogen, and all other reagents were Sigma-Aldrich products unless specially noted.

### Expression and purification of recombinant proteins

Protein expression was induced overnight in *E. coli* BL21 (DE3) strain at 22°C with 0.4 mM isopropyl-β-D-thiogalactopyranoside (IPTG) when OD600 reached 0.8∼1.0. Affinity purification of GST-DDs was performed by using glutathione sepharose (GE Healthcare, USA), and that of LFN-NleB, SseK1, SseK2, and SseK3 and His-NleB, SseK1, SseK2, and SseK3 was conducted by using Ni^2+^-charged column chromatography (GE Healthcare, USA), following the manufactures’ instructions. GST-DDs were further purified by ion exchange chromatography. All the purified recombinant proteins were concentrated and stored in the buffer containing 20 mM Tris-HCl (pH 8.0), 150 mM NaCl, and 5 mM dithiothreitol and protein purity was examined by SDS-PAGE, followed by Coomassie Blue staining.

### Cell culture and NF-κB luciferase reporter assay

HEK293T cells and HeLa cells obtained from the American Type Culture Collection (ATCC) and MEF cells provided by S. Ghosh were grown in DMEM (GIBCO) medium supplemented with 10% FBS (GIBCO and BI), 2 mM L-glutamine (GIBCO), 100 U/ml penicillin, and 100 mg/ml streptomycin (GIBCO). These cells were cultivated at 37°C in the presence of 5% CO_2_. Vigofect (Vigorus) was used for HEK293T cell transfection and jetPRIME (PolyPlus) for HeLa cell transfection, following the respective manufacturer’s instructions. Luciferase activity was determined 24 h after transfection by using the dual luciferase assay kit (Promega) according to the manufacturer’s instructions.

### Immuofluorescence

For immunofluorescence staining, cells were fixed with 4% paraformaldehyde (PFA) for 20 min at room temperature and permeabilized for 10 min with 0.5% Triton X-100 in PBS. Then cells were blocked at room temperature with 5% BSA for 1 hour, followed by the incubation with the indicated primary antibody and secondary antibody. All the cell nuclei were counterstained with DAPI before imaging.

### Immunoprecipitation

The 293T cells seeded in 6-well plates at a confluency of 60-80% were transfected with a total of 5 μg plasmids. Twenty-four hours after transfection, cells were washed in 1×PBS and lysed in lysis buffer A containing 25 mM Tris-HCl, pH 7.6, 150 mM NaCl, 10% glycerol, and 1% Triton, supplemented with a protease inhibitor cocktail (Roche). Cells were collected and centrifuged under 13,200 rpm at 4°C for 15 min. Pre-cleared lysates were subjected to anti-Flag M2 immunoprecipitation, following the manufacturer’s instructions. The beads were washed four times with lysis buffer B containing 25 mM Tris-HCl, pH 7.6, 150 mM NaCl, 10% glycerol, and 0.5% Triton, and the immunoprecipitates were eluted by 1×SDS sample buffer, followed by standard immunoblotting analysis. All the immunoprecipitation assays were performed for more than three times and representative results are shown in figures.

### Bacterial infection of mammalian cells

The infection was performed as described previously(20, 55). Briefly, HeLa cells or MEF cells were seeded in 96 well plates at a concentration of 2×10^4^ per well one day before infection. For EPEC and *C. rodentium* infection, a single colony was incubated overnight in a static LB culture at 37°C. Bacterial strains were then diluted by 1:40 in antibiotic-free DMEM supplemented with 1 mM IPTG and cultured at 37°C in the presence of 5% CO_2_ for an additional 4 h to OD600 approximately at 0.4∼0.6. For *Salmonella* infection, a single colony was incubated in LB culture with shaking at 37°C overnight. The infection strains were diluted by 1:33 in LB and cultured at 37°C with shaking for another 3 h. Infection was performed at a multiplicity of infection (MOI) of 200 in the presence of 1 mM IPTG for EPEC and *C. rodentium,* or at MOI of 100 for *Salmonella,* with a centrifugation at 800 g for 10 minutes at room temperature to promote infection. Infected cells were incubated at 37°C, in the presence of 5% CO_2_ for 2 hours. Then cells were washed four times with PBS and the extra bacteria were killed with 200 μg/ml gentamicin for EPEC and *C. rodentium,* or with 100 μg/ml gentamicin without penicillin and streptomycin for *Salmonella*. One-hour CHX pretreatment (1 μg/ml or 2 μg/ml) was performed to sensitize TRAIL (200 ng/ml)- or TNF-α (20 ng/ml)-stimulated cell death. Cell survival was then determined 15 h after treatment with TRAIL and TNF-α by using the CellTiter-Glo® Luminescent Cell Viability Assay kit (Promega).

### Preparation of arginine-GlcNAc beads and enrichment of auto-arginine-glycosylated proteins

All the operations in this experiment were carried out at 4°C. The 30 μl protein A/G plus agarose beads (Santa Cruz) were washed three times with 1 ml of protein-dissolved buffer (50 mM Tris-HCl, pH 7.4) at 6000 g for 1 minute, then incubated overnight with 3 μg of mouse anti-Arg-GlcNAc antibody. The coupled anti-Arg-GlcNAc beads were harvested at 1000 g for 5 minutes and washed with 1 ml of protein-dissolved buffer for three times. Well-prepared beads should be used immediately. The 50 μg of purified proteins in protein-dissolved buffer were then added to the prepared anti-Arg-GlcNAc beads, followed by 4-hour incubation. After incubation, the antibody beads were centrifuged at 3000 g for 5 minutes and washed three times with 1 ml of protein-dissolved buffer. Beads enriched with auto-arginine-glycosylated proteins were used *in vitro* glycosylation assay.

### *In vitro* glycosylation assay

Recombinant protein LFN-NleB purified from bacterial cultures was enriched using methods mentioned above. Flag-FADD expressed in 293T cells was immunopurified and immobilized on the Flag M2 beads. The beads were then incubated with 5 μg NleB protein and the prepared auto-arginine-glycosylated beads for 2 h at 37°C in 40 μl buffer containing 25 mM HEPES, pH 7.5, 2 mM MnCl_2_, 0.2 mM UDP-GlcNAc (Perkin Elmer), and 1 mM DTT, with other proper controls. To detect *in vitro* glycosylation, the reaction mixtures were separated with SDS-PAGE gels, followed by western blot.

### Liquid chromatography-mass spectrometry analysis

The 10 μg of intact recombinant protein was injected and separated by reversed-phase liquid chromatography in a Dionex Ultimate 3000 HPLC system (Thermo Scientific, USA) using a C4 capillary column (MAbPacTM RP, 4 μm, 2.1 × 50 mm, Thermo Scientific, USA) at a flow rate of 0.3 ml/min with buffer A containing 0.1% Formic acid (Thermo Scientific, USA) and buffer B with a linear 10 min gradient from 5% to 100%, containing 0.1% Formic acid, 80% acetonitrile (Merck, Germany). The eluted proteins were sprayed into a Q Exactive Plus mass spectrometer (Thermo Scientific, USA) equipped with a Heated Electrospray Ionization (HESI-II) Probe (Thermo Scientific, USA). Mass accuracy of the intact protein was analyzed by using the Thermo Scientific Protein Deconvolution program.

### Tryptic digest of gel-separated proteins

Samples were separated by SDS-PAGE, visualized with Coomassie G-250 or by ProteoSilverä PlusSilver Stain Kit (SIGMA, USA) according to protocol instructions. Bands were excised and destained in 50% acetonitrile (ACN) and 50 mM NH_4_HCO_3_ solution with shaking at room temperature for 1 h. Destained samples were then washed with the buffer containing 300 μl 100% ACN, and the buffer was removed after 10 min incubation. Subsequently, the resultant samples were speed-vacuumed for at least 10 min. Rehydrated samples were then incubated with the reducing buffer containing 300 μl DTT solution (10 mM DTT in 100mM NH_4_HCO_3_) for 1 h at 56°C with shaking. The reducing buffer was then removed and the reduced samples were washed twice with 100% ACN, each for 10 minutes to remove residual DTT. Reduced samples were subsequently alkylated with 60 mM Iodoacetamide (IAM) in 50 mM NH_4_HCO_3_ in the dark for 30 minutes at room temperature. Alkylated samples were hydrated with 100% ACN for 10 min incubation. Reduced and alkylated samples were then digested with enough trypsin for 16 h at 37°C . Trypsin was removed and the digested peptides were collected by extraction buffer (50% ACN, 5% FA). Peptides were desalted using C18 stage tips and dried by concentrator at 60°C and analyzed by a Q Exactive Plus mass spectrometer.

### HCD mass spectrometry analysis of tryptic peptides

The digested peptide samples were analyzed by using a Q Exactive Plus mass spectrometer and an EASY nano Liquid chromatography (EASY nLC 1200, Thermo Scientific) with an EASY nano electrospray interface. The nano liquid chromatography system was equipped with a Thermo Scientific Acclaim Pepmap nano-trap column (C18, 5 μm, 100 Å, 100 μm × 2 cm) and a Thermo Scientific EASY-Spray column (Pepmap RSLC, C18, 2 μm, 100 Å, 50 μm × 15 cm). The nano liquid chromatography used solvent A (0.1% Formic acid) and solvent B (80% CH_3_CN/0.1% formic acid), whose gradients were as follows: 0∼8% B for 3 min, 8∼28% B for 42 min, 28∼38% B for 5 min, 38∼100% B for 10 min. The mass spectra were searched against the UniProt database, and were analyzed by using the Thermo Scientific Proteome Discoverer 2.2 program.

Mice infection and *C. rodentium* colonization assays

All animal experiments were conducted following the Chinese National Ministry of Health guidelines for housing and care of laboratory animals, and the experiments were performed in accordance with institutional regulations made by the Institutional Animal Care and Use Committee at Taihe Hospital, Hubei University of Medicine. The C57BL/6 male mice 5-6 weeks old were maintained in the specific pathogen-free environment. All the mice were randomly divided into each experimental group with no blind mice, and were housed individually in high efficiency particulate air (HEPA)-filtered cages with sterile bedding. Independent experiments were performed using 6-8 mice per group. The mice infection was performed as described previously(20, 22). Briefly, *nleB* was deleted from *C. rodentium* strain ICC168 by standard homologous recombination using the suicide vector pCVD442. Before oral inoculation and harvesting, *C. rodentium* WT strain and the indicated derivatives were prepared in LB broth by overnight bacterial culture with shaking at 37°C . Mice were orally inoculated with 200 μl bacteria containing approximately 2×10^9^ CFU in PBS via a gavage needle. The number of viable bacteria used as the inoculum was determined by plating onto the LB agar containing the appropriate antibiotics. Stool samples were harvested and recovered aseptically at various time points after inoculation, and the number of viable bacteria per gram of stool was determined after homogenization in PBS, and the homogenized stool samples were plated onto LB agar containing the appropriate antibiotics. Eight days after inoculation, colons were collected aseptically, weighed, and homogenized in PBS. Homogenates were serially diluted and plated to determine the CFU counts. Colonization data were analyzed using student’s T test in the software GraphPad Prism. *P* < 0.05 was considered significant.

### Statistical analysis

All the values of at least three independent experiments are presented. Statistical analysis was performed using Student’s T test in the commercial software GraphPad Prism to compare two experimental groups. The comparison of multiple groups was conducted by using one-way analysis of variance (ANOVA). *P* < 0.05 was considered significant.

## Acknowledgments

We thank Jun Lv and Jin Yang for preparing gene knockout bacteria strains. We also thank members of the Li laboratory and the central laboratory of Taihe hospital for helpful discussions and technical assistance.

## Author Contributions

**Conceptualization:** Juan Xue, Xing Pan, Lijie Du, Shan Li

**Funding acquisition:** Shan Li

**Investigation:** Juan Xue, Xing Pan, Lijie Du, X.Z., X.C.

**Supervision:** Shan Li

**Visualization:** Juan Xue, Xing Pan, Lijie Du, Shan Li

**Writing-original draft:** Juan Xue, Xing Pan, Shan Li

**Writing-review&editing:** Juan Xue, Shan Li

## Supporting information

## Funding

This work was supported by the National Key Research and Development Programs of China 2018YFA0508000, Fundamental Research Funds for the Central Universities 2662017PY011, 2662018PY028, 2662019YJ014, 2662018JC001and Huazhong Agricultural University Scientific & Technological Self-Innovation Foundation 2017RC003 to S. Li.

**S1 Fig.**
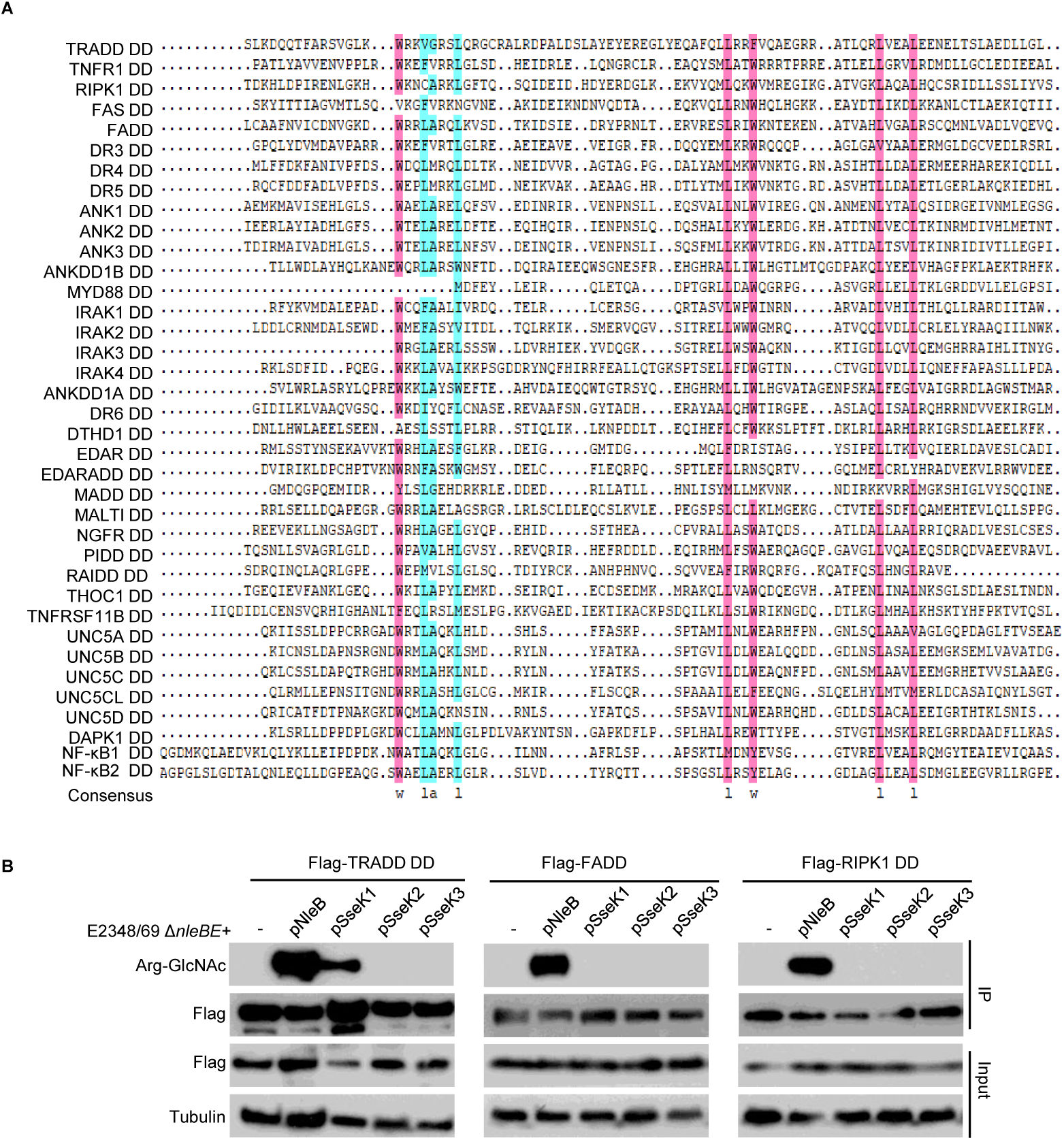
NleB hijacks death receptor signaling via TRADD, FADD, and RIPK1 in EPEC infection system, SseK1 hijacks via TRADD. (**A**) Multiple sequence alignment of 37 death domains (DDs) from human death domain-containing proteins. (**B**) Identification of the physiological substrates of arginine GlcNAc transferase NleB/SseKs during EPEC infection of mammalian cells. 293T cells transfected with pCS2-1Flag-TRADD DD, pCS2-1Flag-FADD DD, and pCS2-1Flag-RIPK1 DD were infected with the indicated modified EPEC strains. After infection, cells were lysed, and proteins were immunoprecipitated with FLAG M2 beads. Samples were loaded onto SDS-PAGE gels and were immunoblotted with anti-Flag, anti-Arg-GlcNAc, and a loading control anti-tubulin. Blot data were derived from at least three independent experiments.

**S2 Fig.**
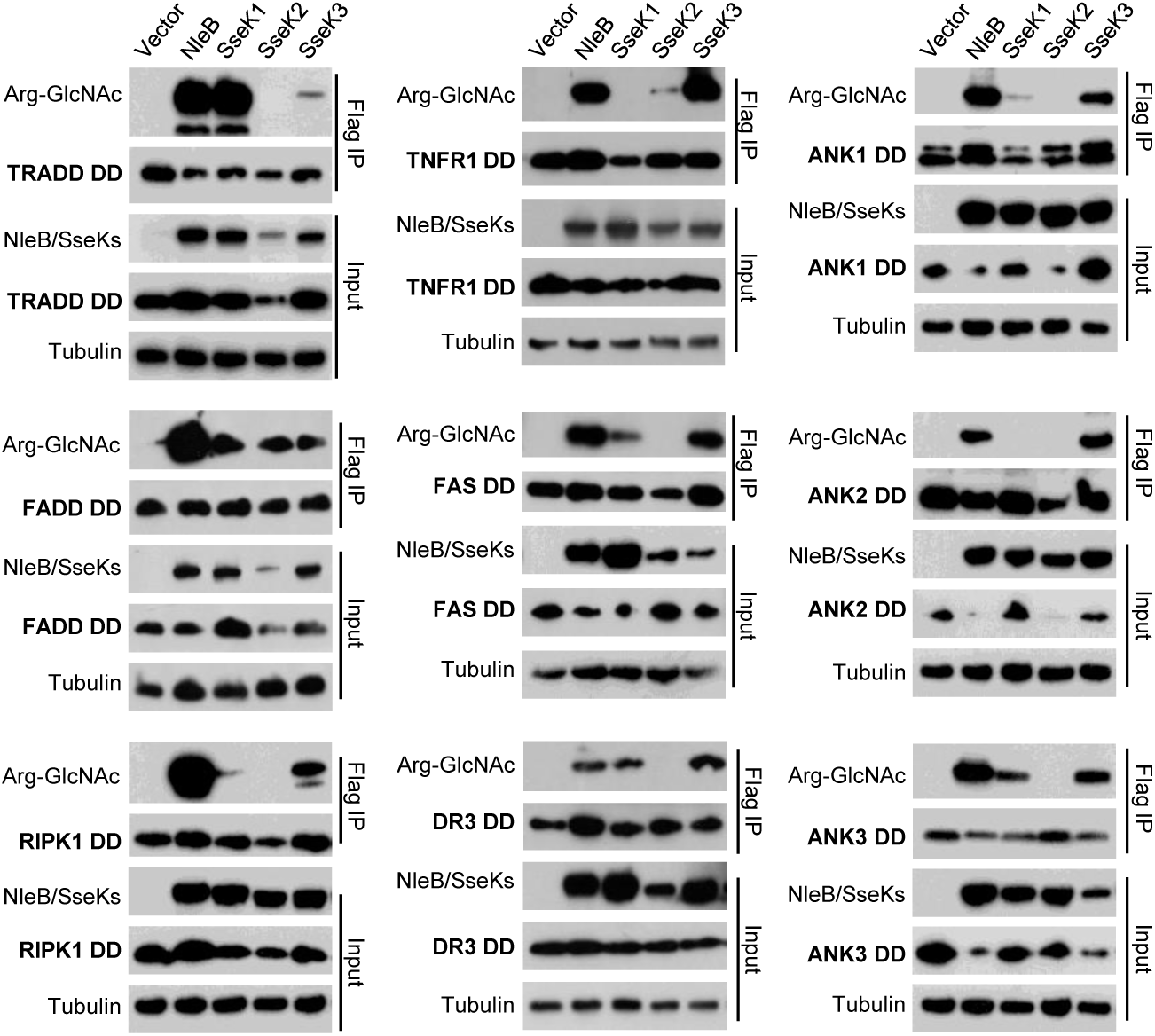
Arginine-GlcNAcylation profile of 12 human death domains containing conserved arginine by NleB/SseKs in overexpression system. 293T cells were transfected with the indicated plasmid combinations. The samples of anti-Flag immunoprecipitates (Flag IP) and total cell lysates (Input) were immunoblotted with corresponding antibodies. Tubulin were used as a loading control. Blot data were derived from at least three independent experiments.

**S3 Fig.**
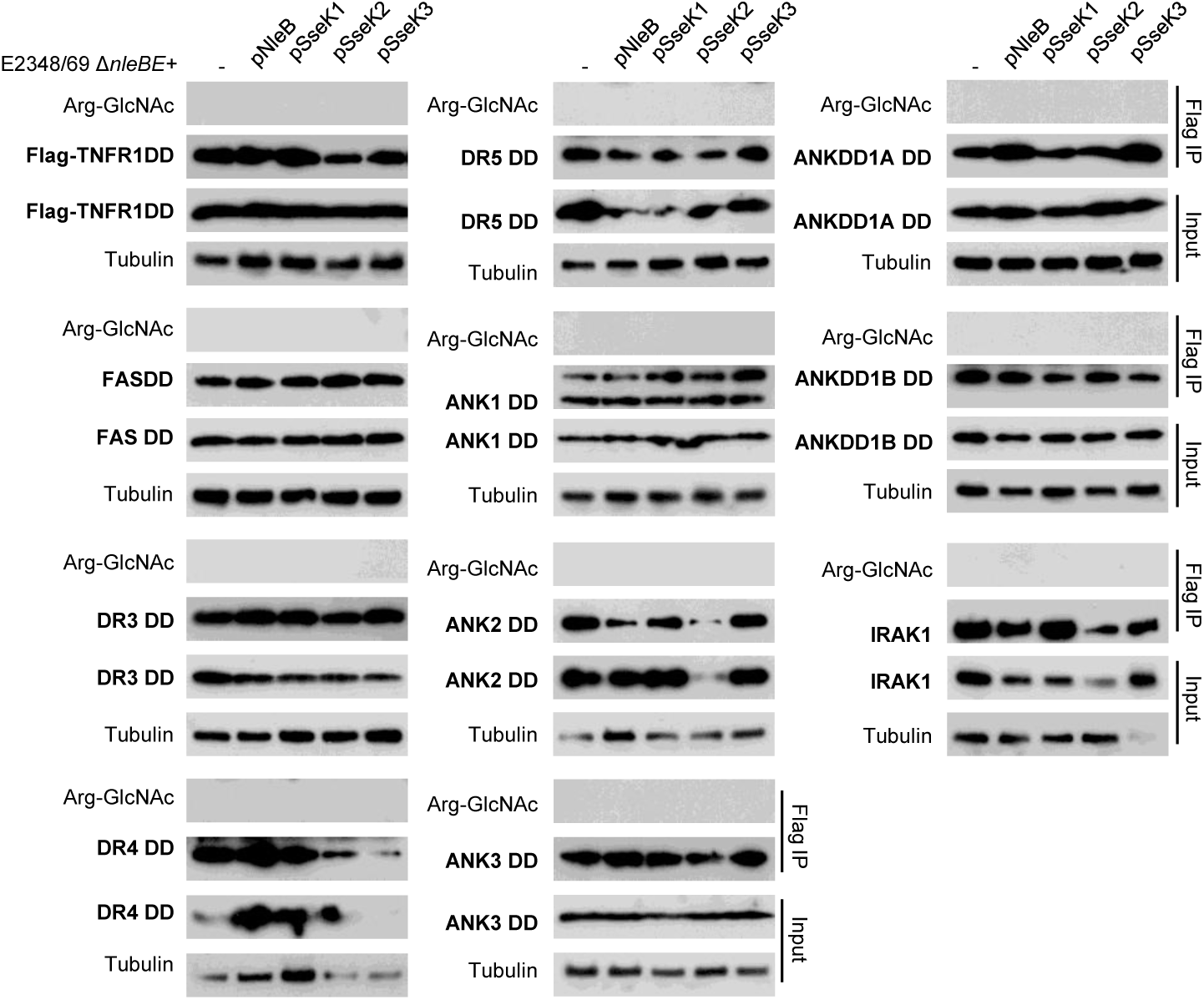
Arginine-GlcNAcylation profile of human death domains containing conserved arginine by NleB/SseKs during EPEC infection. 293T cells expressing the indicated death domains were infected with some modified EPEC mutants which was complemented with plasmids expressing wild type NleB/SseKs. After infection, cells were lysed and proteins were immunoprecipitated with FLAG M2 beads. Samples were loaded onto SDS-PAGE gels and were immunoblotted with anti-Flag, anti-Arg-GlcNAc, and a loading control anti-tubulin. Blot data were derived from at least three independent experiments.

**S4 Fig.**
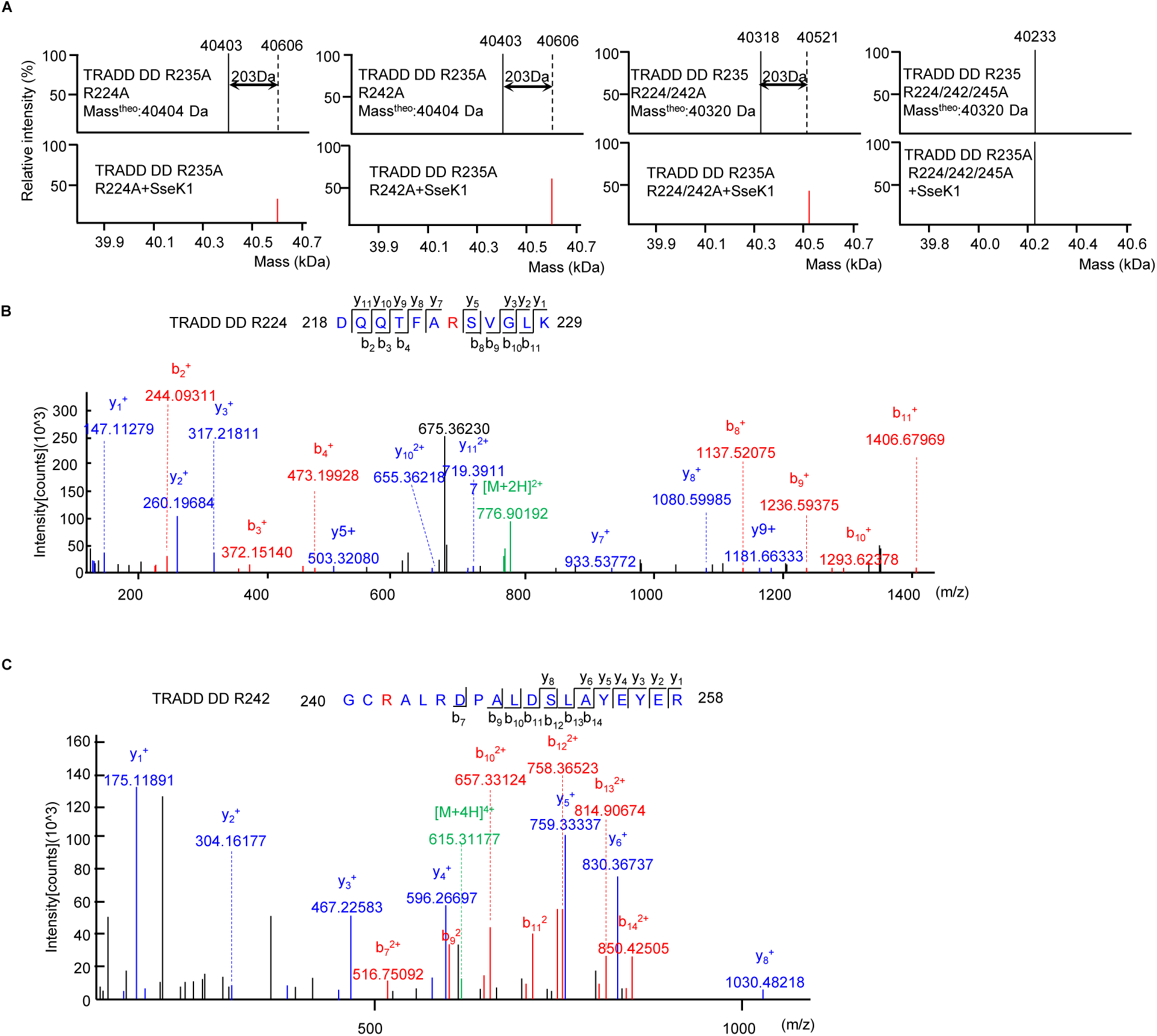
SseK1 glycosylates TRADD at R224 and R242. **(A)** ESI-MS determination of the total mass of site-directed TRADD mutants purified from bacteria. The resulting mass spectra were shown. The black bar and red bar denote unmodified and GlcNAcylated TRADD DD, respectively. (**B** and **C**) HCD spectra showing glycosylation of TRADD DDR235A at Arg224 (**B**) and Arg242 (**C**). The fragmentation patterns of the generated b and y ions were exhibited along the peptide sequence on top of the spectrum.

**S5 Fig.**
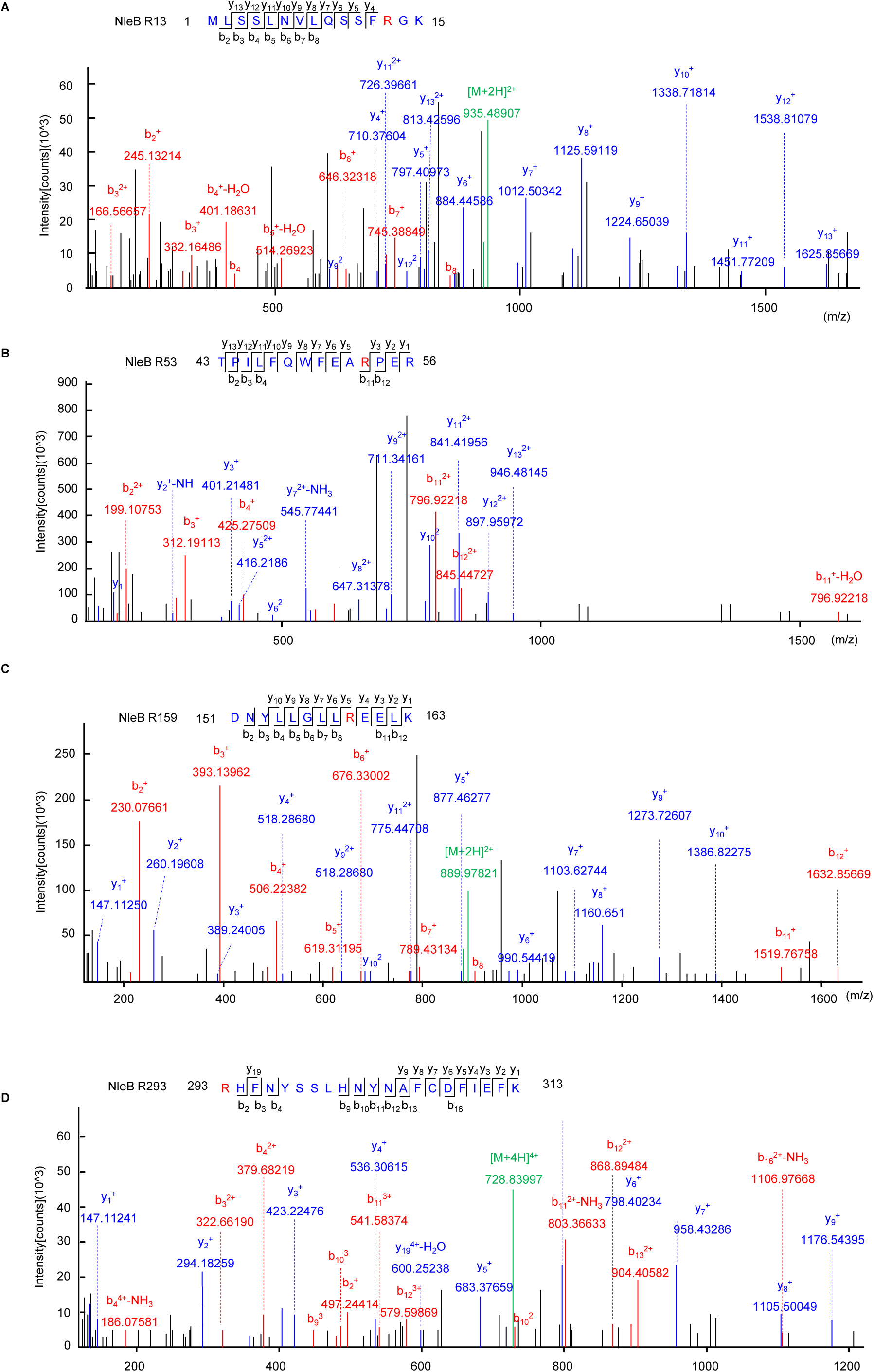
HCD analysis of the peptides GlcNAclated by NleB. HCD mass spectrum of Arg13 (A), Arg53 (B), Arg159 (C), and Arg293 (D) containing tryptic peptide from NleB in bacteria. The fragmentation patterns of the generated b and y ions were displayed along the peptide sequence on top of the spectrum.

**S6 Fig.**
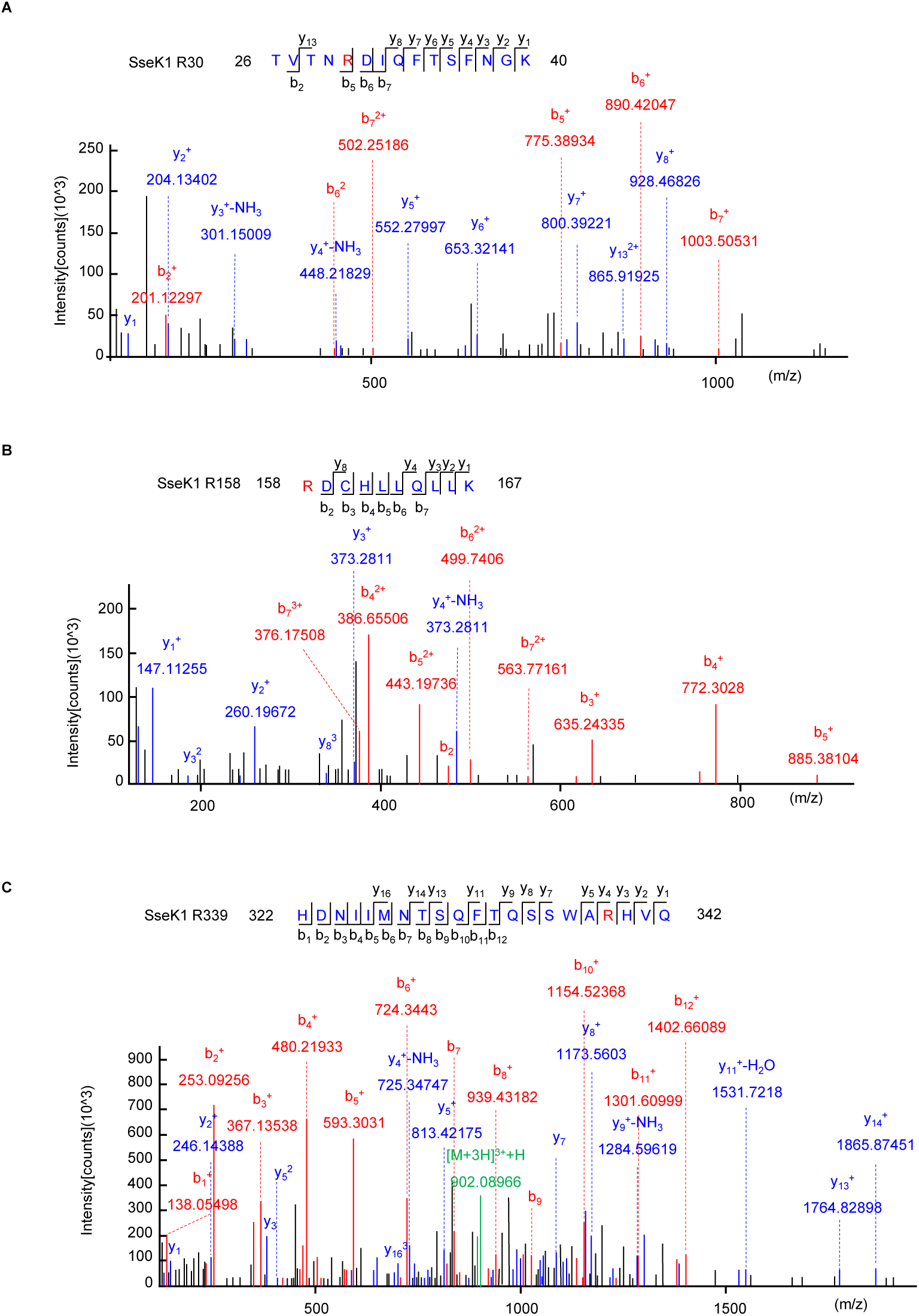
HCD analysis of the peptides GlcNAclated by SseK1. HCD mass spectrum of Arg30 (**A**), Arg158 (**B**), and Arg339 (**C**) containing tryptic peptide from SseK1 in bacteria. The fragmentation patterns of the generated b and y ions were shown along the peptide sequence on top of the spectrum.

**Figure S7.**
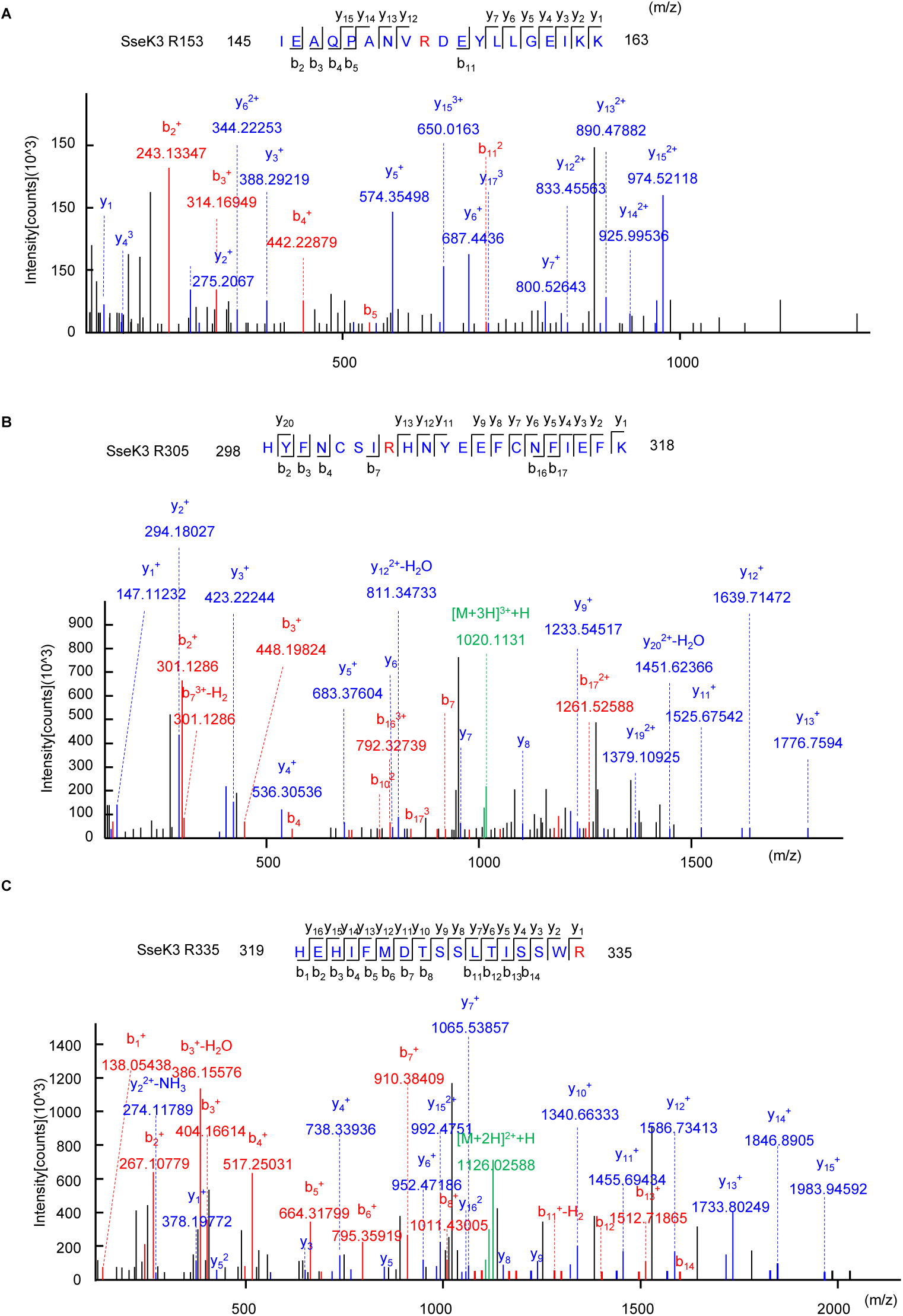
HCD analysis of the peptides GlcNAclated by SseK3. HCD mass spectrum of Arg153 (**A**), Arg305 **(B**), and Arg335 (**C**) containing tryptic peptide from Ssek3 in bacteria. The fragmentation patterns of the generated b and y ions were exhibited along the peptide sequence on top of the spectrum.

**S8 Fig.**
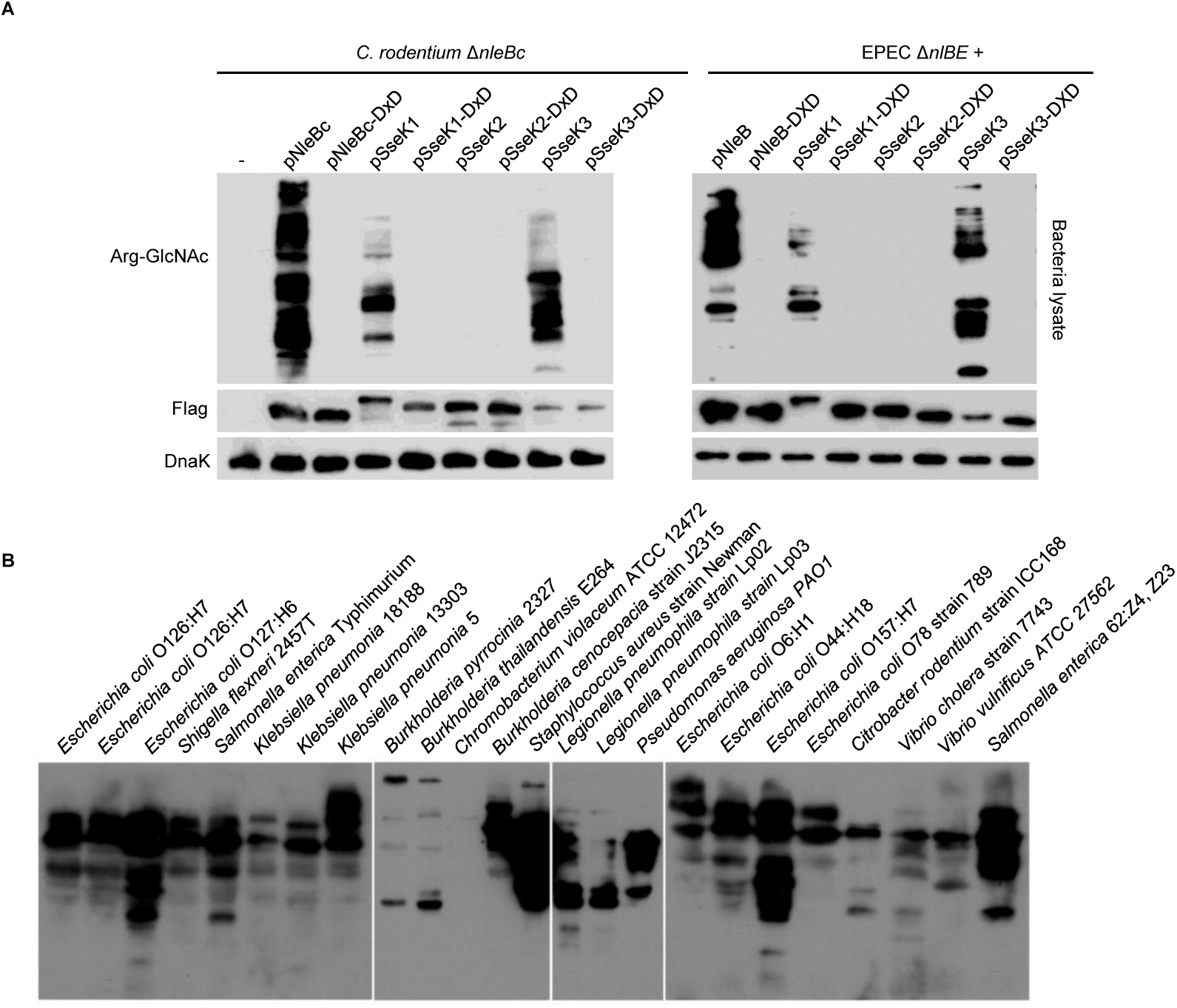
Arginine-GlcNAcylation widely exists in various bacteria. (**A**) Different arginine-GlcNAcylation protein patterns in C. rodentium and EPEC strains complemented with NleB/SseKs and their enzyme inactivated mutants. (**B**) Analysis of arginine-GlcNAcylation in various bacteria.

**S1 Table.** EPEC strains used in this study.

| EPEC Strains |  |  |
| --- | --- | --- |
| Name | Relevant genotype and features | Reference |
| E2348/69 | Wild type EPEC O127:H6 | (20) |
| EPEC $\Delta escN$ | Lacking T3SS | (20) |
| SC3909 | E2348/69 $\Delta nleBE$ | (20) |
| SC3909+pNleB | SC3909+pTRC99A-NleB | (20) |
| SC3909+pNleB-DxD | SC3909+pTRC99A-NleB-DxD | (20) |
| SC3909+pSseK1 | SC3909+pTRC99A-SseK1 | This study |
| SC3909+pSseK1-DxD | SC3909+pTRC99A-SseK1-DxD | This study |
| SC3909+pSseK2 | SC3909+pTRC99A-SseK2 | This study |
| SC3909+pSseK2-DxD | SC3909+pTRC99A-SseK2-DxD | This study |
| SC3909+pSseK3 | SC3909+pTRC99A-SseK3 | This study |
| SC3909+pSseK3-DxD | SC3909+pTRC99A-SseK3-DxD | This study |
| SC3909+pNleB-Flag | SC3909+pTRC99A-NleB-Flag | This study |
| SC3909+pNleB-DxD-Flag | SC3909+pTRC99A-NleB-DxD-Flag | This study |
| SC3909+pSseK1-Flag | SC3909+pTRC99A-SseK1-Flag | This study |
| SC3909+pSseK1-DxD-Flag | SC3909+pTRC99A-SseK1-DxD-Flag | This study |
| SC3909+pSseK2-Flag | SC3909+pTRC99A-SseK2-Flag | This study |
| SC3909Δ+pSseK2-DxD-Flag | SC3909+pTRC99A-SseK2-DxD-Flag | This study |
| SC3909+pSseK3-Flag | SC3909+pTRC99A-SseK3-Flag | This study |
| SC3909+pSseK3-DxD-Flag | SC3909+pTRC99A-SseK3-DxD-Flag | This study |
| SC3909+pNleB-HA | SC3909+pTRC99A-NleB-HA | This study |
| SC3909+pNleB-DxD-HA | SC3909+pTRC99A-NleB-DxD-HA | This study |
| SC3909+pSseK1-HA | SC3909+pTRC99A-SseK1-HA | This study |
| SC3909+pSseK1-DxD-HA | SC3909+pTRC99A-SseK1-DxD-HA | This study |
| SC3909+pSseK2-HA | SC3909+pTRC99A-SseK2-HA | This study |
| SC3909Δ+SseK2-DxD-HA | SC3909+pTRC99A-SseK2-DxD-HA | This study |
| SC3909+pSseK3-HA | SC3909+pTRC99A-SseK3-HA | This study |
| SC3909+pSseK3-DxD-HA | SC3909+pTRC99A-SseK3-DxD-HA | This study |
| $\Delta escN$+pNleB-HA | Lacking T3SS, complementary with<br>pTRC99A-NleB(EPEC)-HA | This study |
| SC3909+pNleB(4RA) | SC3909+pTRC99A-NleBR13/53/159/293A-HA | This study |

**S2 Table.** *Salmonella* strains used in this study.

| **Name** | **Reference** |
| --- | --- |
| *Salmonella enterica* serovar Typhimurium SL1344 | This study |
| SL1344+Vector | This study |
| SL1344 $\Delta_{sseK1/2/3}$+Vector | This study |
| SL1344 $\Delta_{sseK1/2/3}$+pSseK1 | This study |
| SL1344 $\Delta_{sseK1/2/3}$+pSseK1(3RA) | This study |
| SL1344 $\Delta_{sseK1/2/3}$+pSseK2 | This study |
| SL1344 $\Delta_{sseK1/2/3}$+pSseK3 | This study |
| SL1344 $\Delta_{sseK1/2/3}$+pSseK3(4RA) | This study |
| SL1344 $\Delta_{sseK1/2/3}$+pSseK3-DxD | This study |

**S3 Table.** Table *C. rodentium* strains used in this study.

| **Name** | **Reference** |
| --- | --- |
| *C. rodentium* ICC168 | (20) |
| CRΔ*nleBc* | (20) |
| Δ*nleBc*+pNleBc | (20) |
| Δ*nleBc*+pNleB-DxD | (20) |
| Δ*nleBc*+pSseK1 | This study |
| Δ*nleBc*+pSseK1-DxD | This study |
| Δ*nleBc*+pSseK2 | This study |
| Δ*nleBc*+pSseK2-DxD | This study |
| CRΔ*nleBc*+pSseK3 | This study |
| Δ*nleBc*+pSseK3-DxD | This study |
| Δ*nleBc*+pNleB-Flag | This study |
| Δ*nleBc*+pNleB-DxD-Flag | This study |
| Δ*nleBc*+pSseK1-Flag | This study |
| Δ*nleBc*+pSseK1-DxD-Flag | This study |
| Δ*nleBc*+pSseK2-Flag | This study |
| Δ*nleBc*+pSseK2-DxD-Flag | This study |
| Δ*nleBc*+pSseK3-Flag | This study |
| Δ*nleBc*+pSseK3-DxD-Flag | This study |
| Δ*nleBc*+pNleB-Flag | This study |
| Δ*nleBc*+pNleB-DxD-Flag | This study |
| Δ*nleBc*+pSseK1-Flag | This study |
| Δ*nleBc*+pSseK1-DxD-Flag | This study |
| Δ*nleBc*+pSseK2-Flag | This study |
| Δ*nleBc*+pSseK2-DxD-Flag | This study |
| Δ*nleBc*+pSseK3-Flag | This study |
| Δ*nleBc*+pSseK3-DxD-Flag | This study |

**S4 Table.**
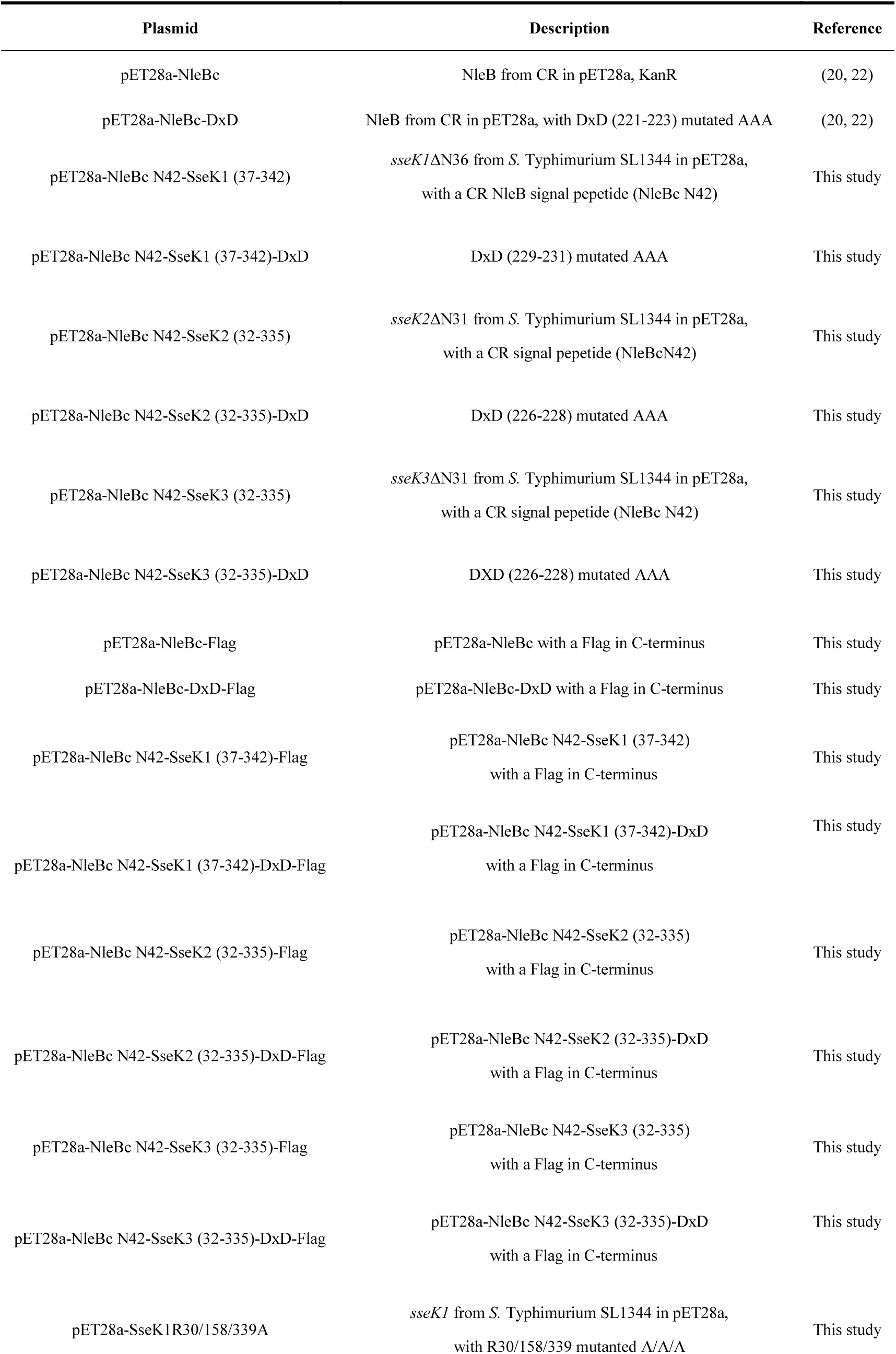
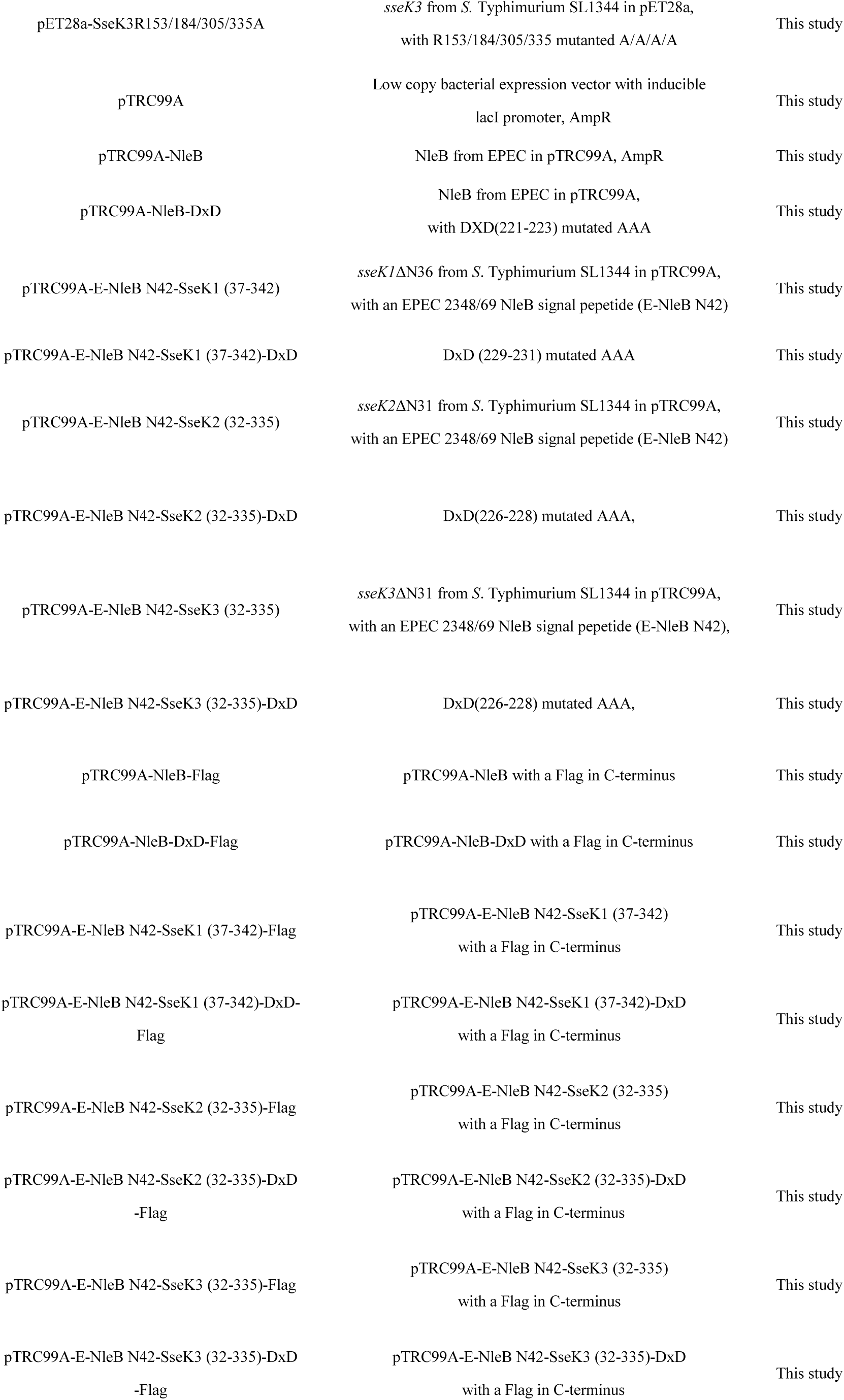
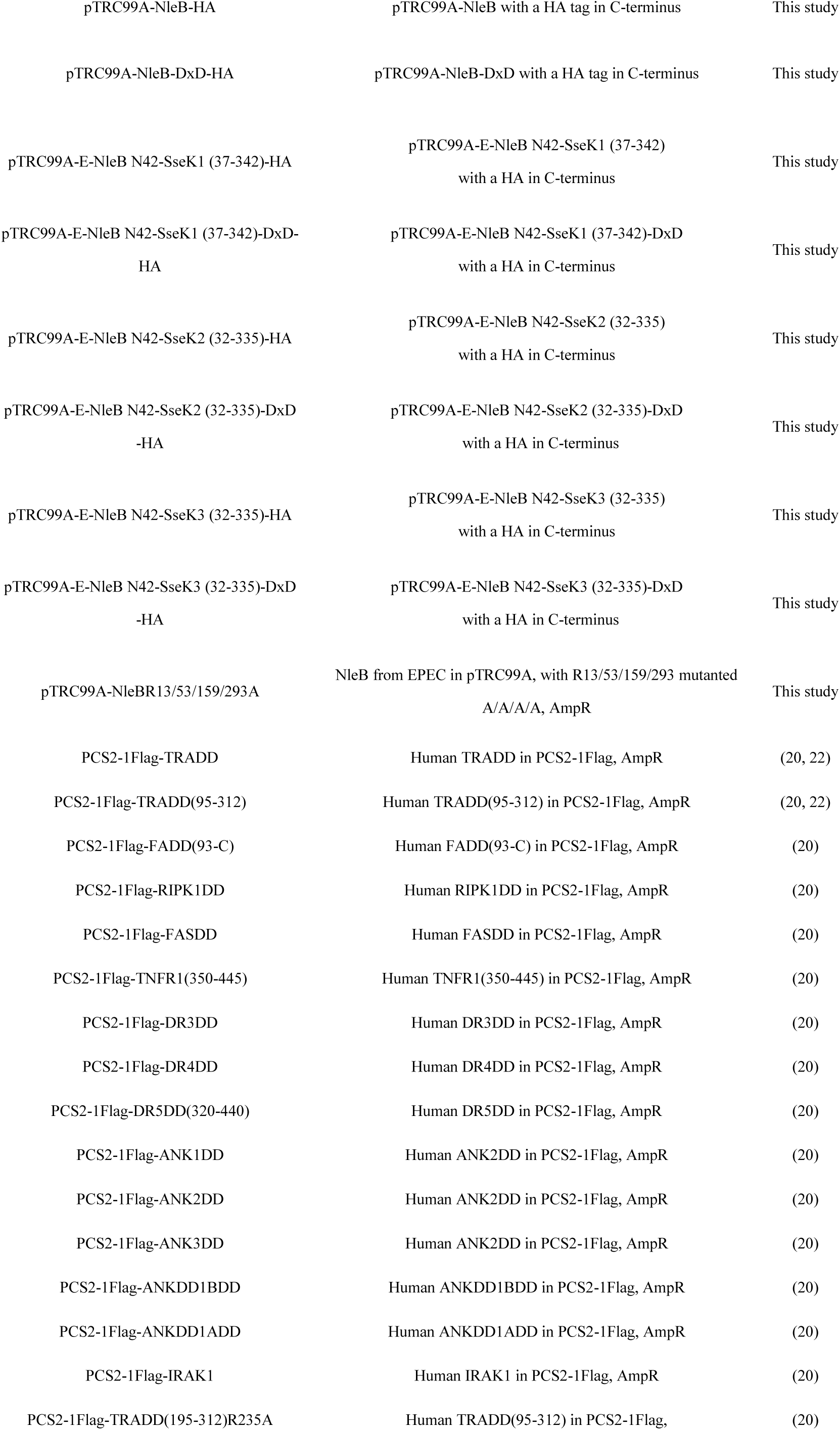
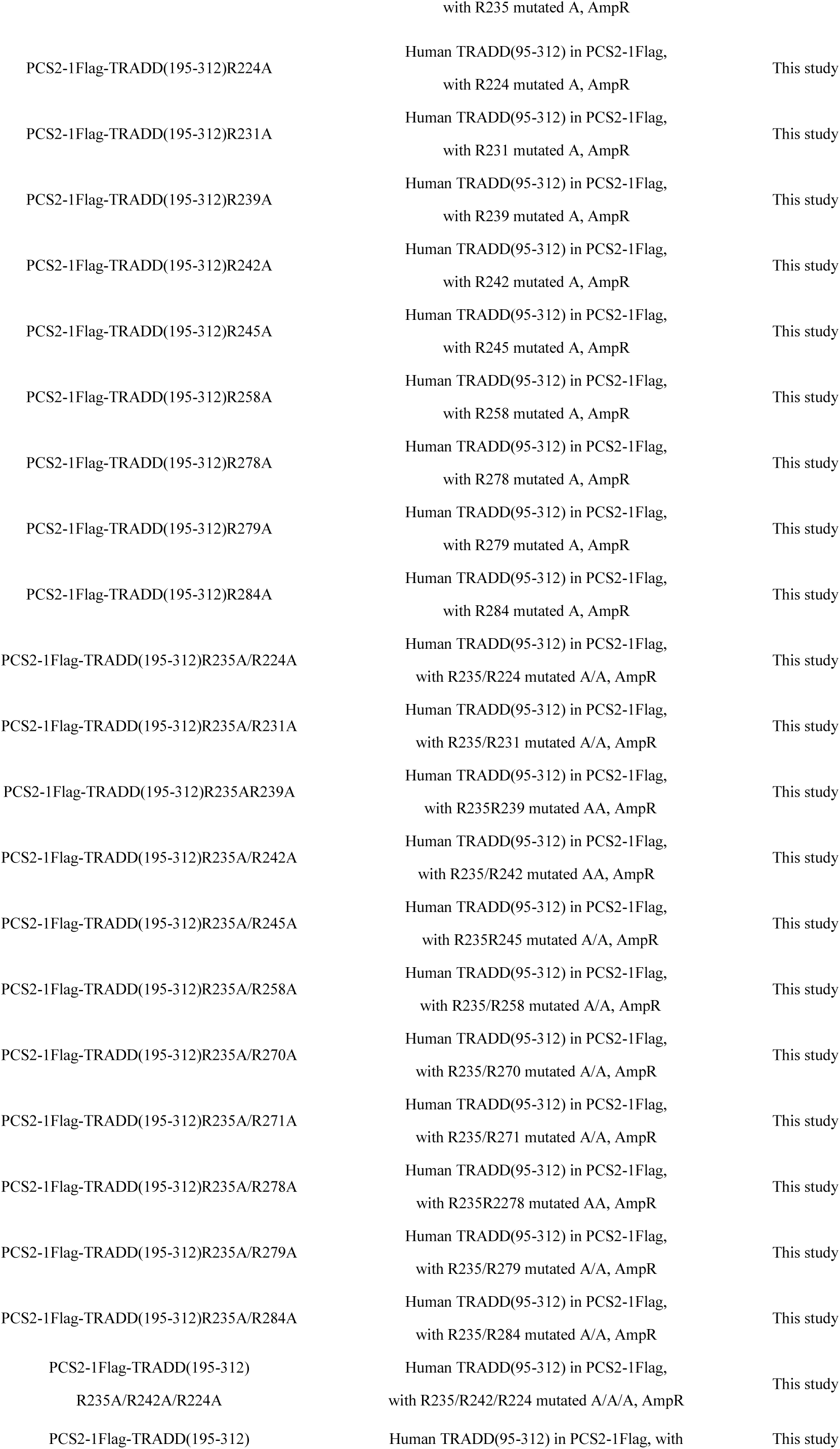
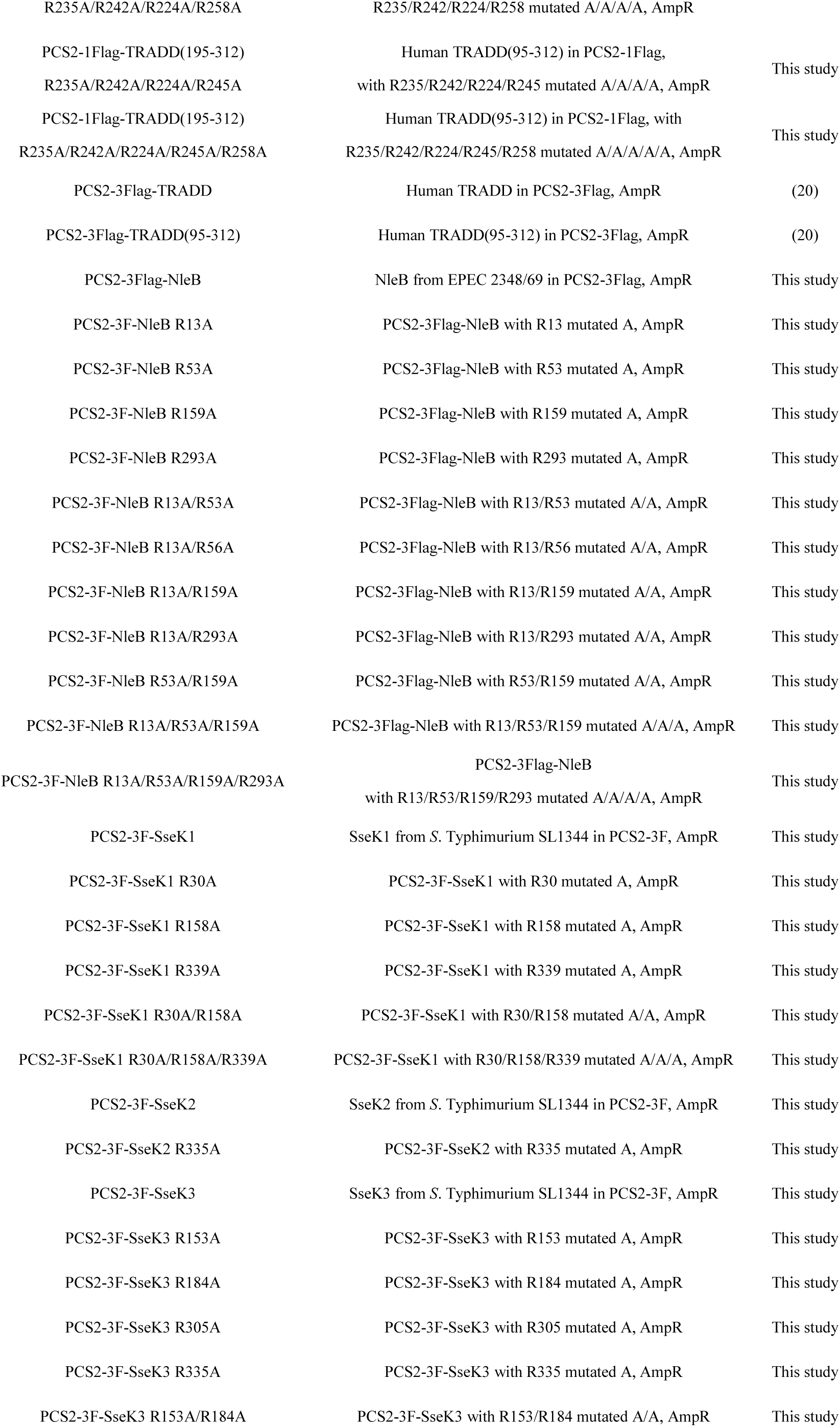
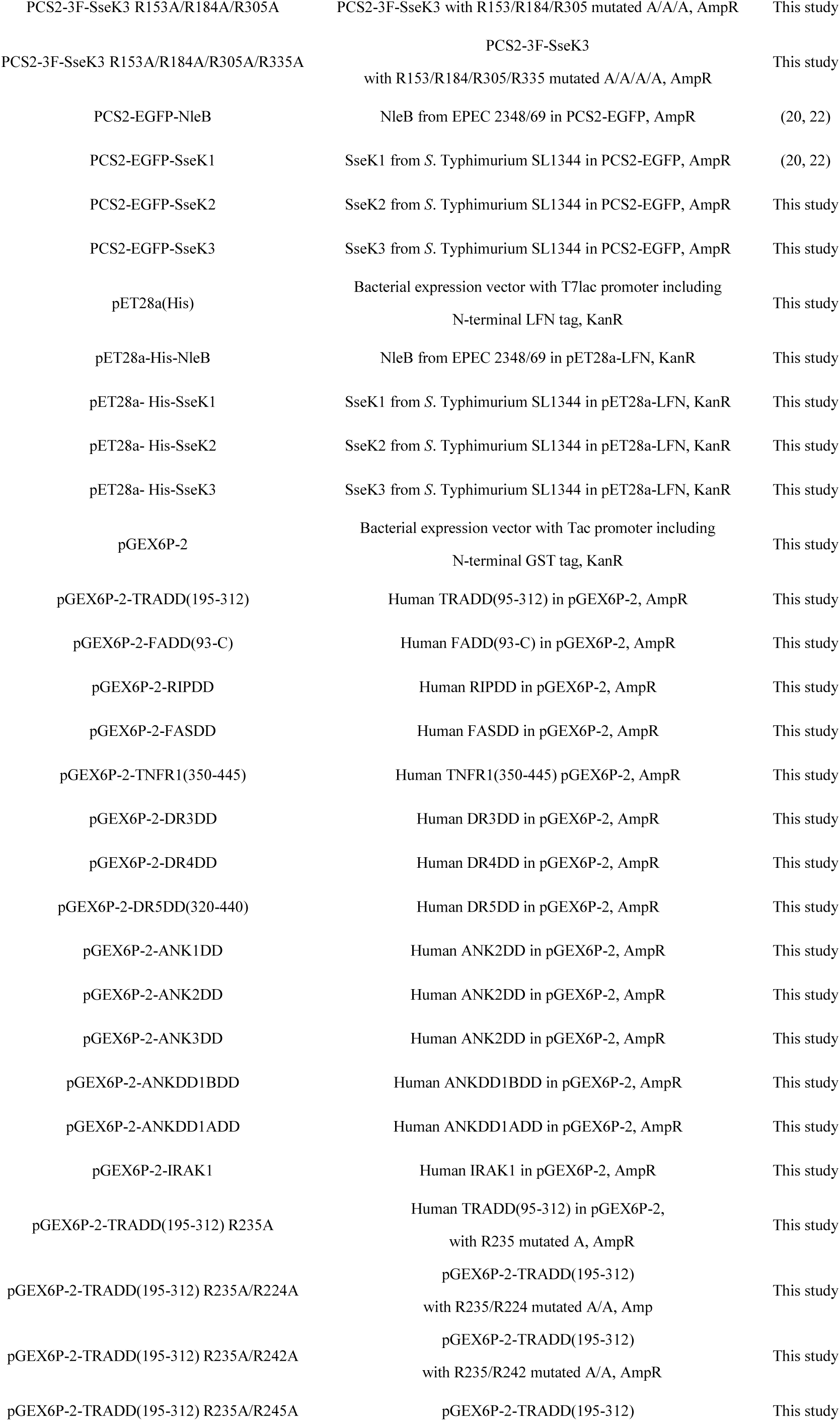
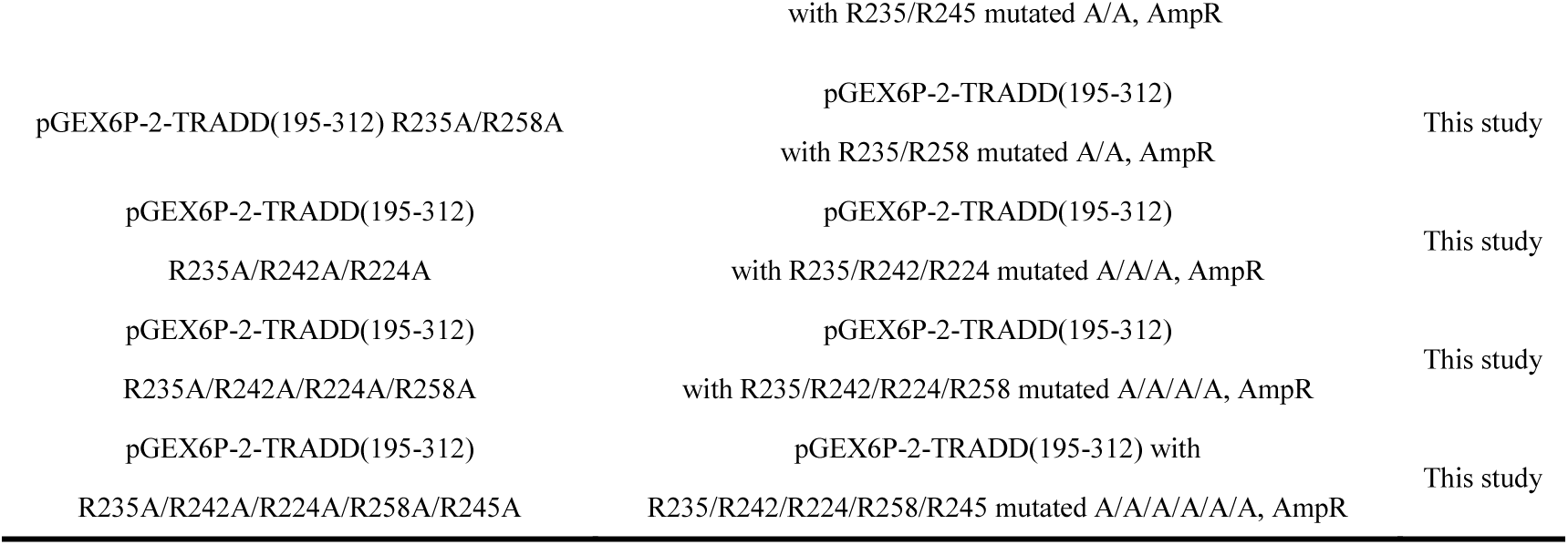
Table Plasmids used in this study.

**S5 Table.**
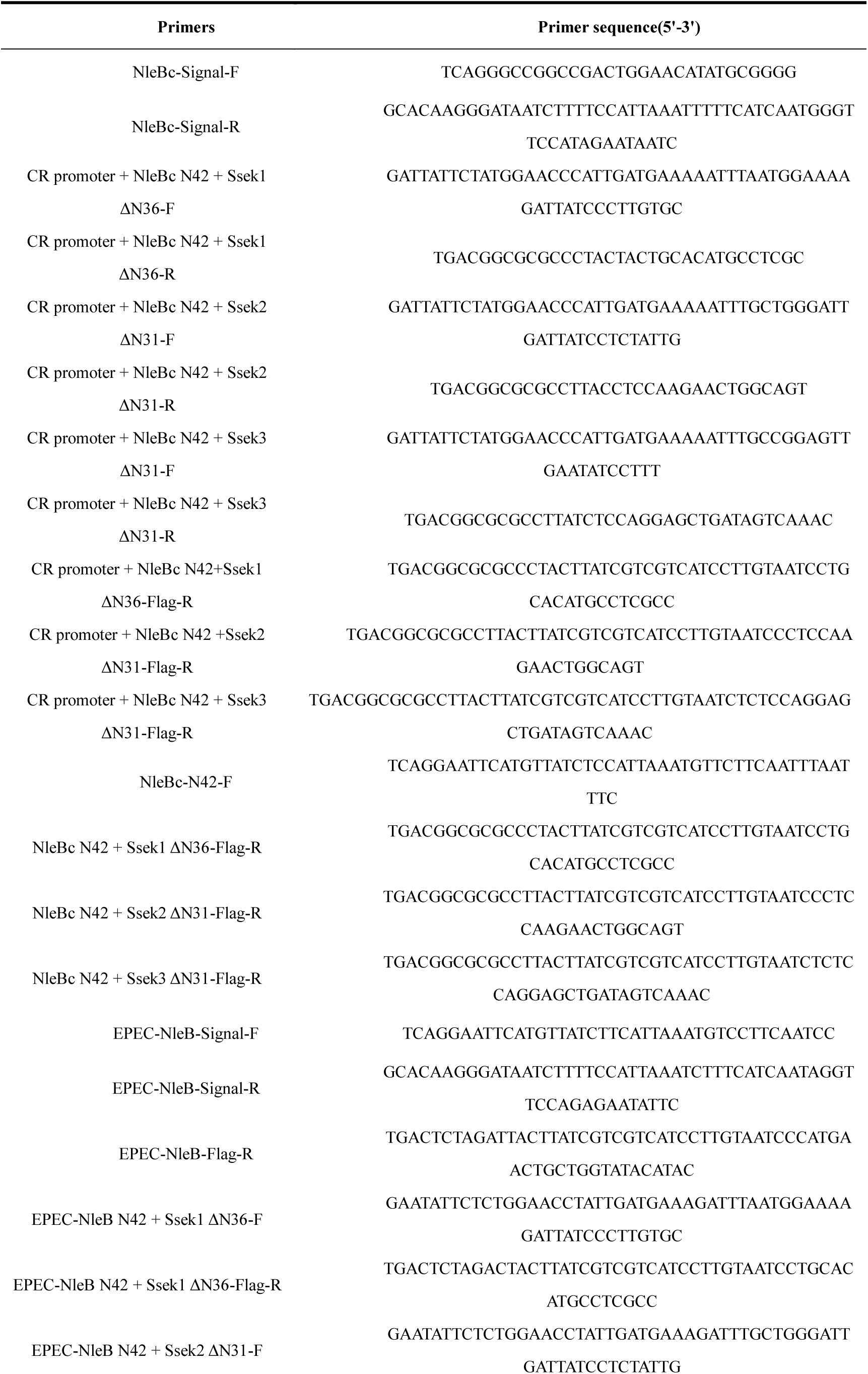
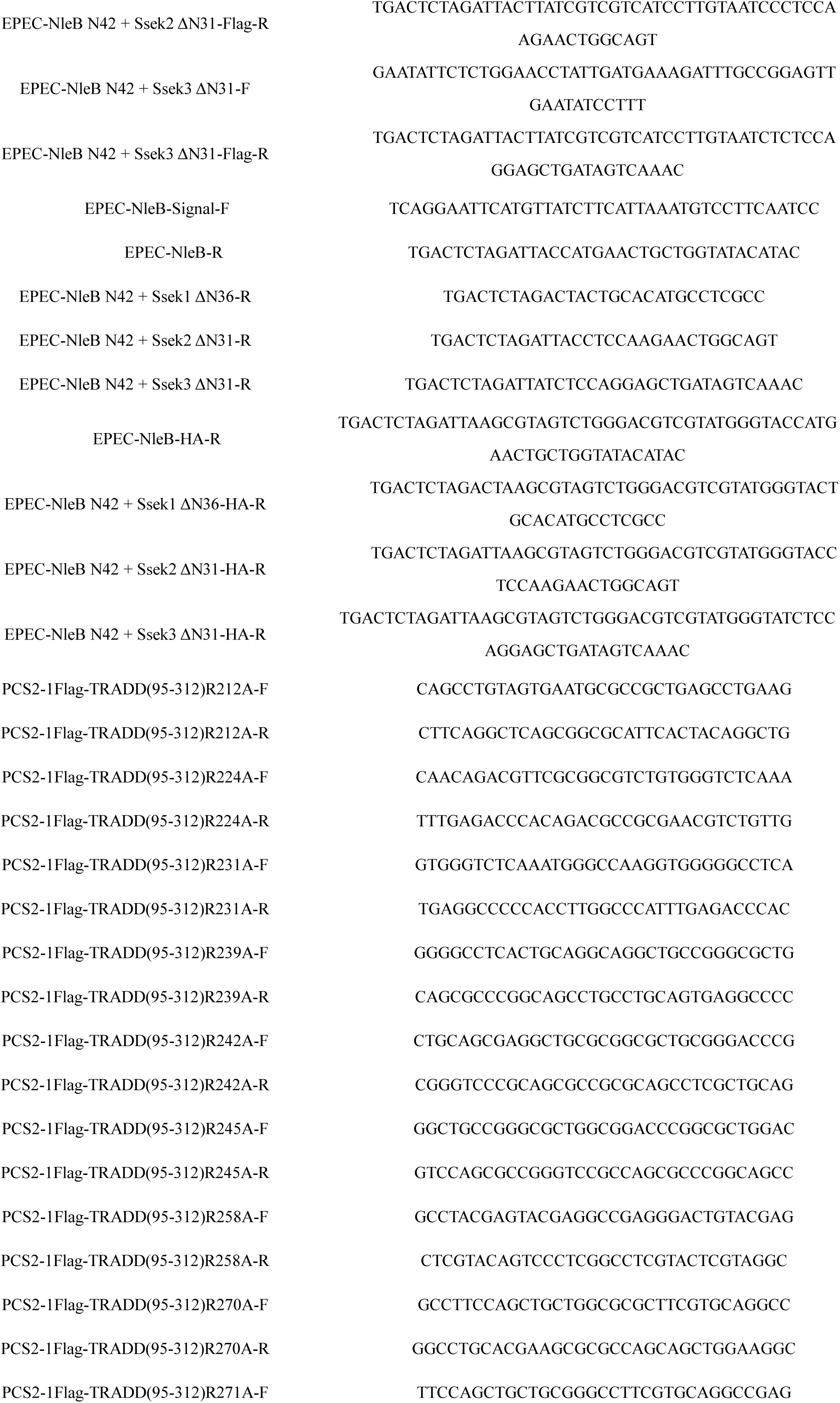
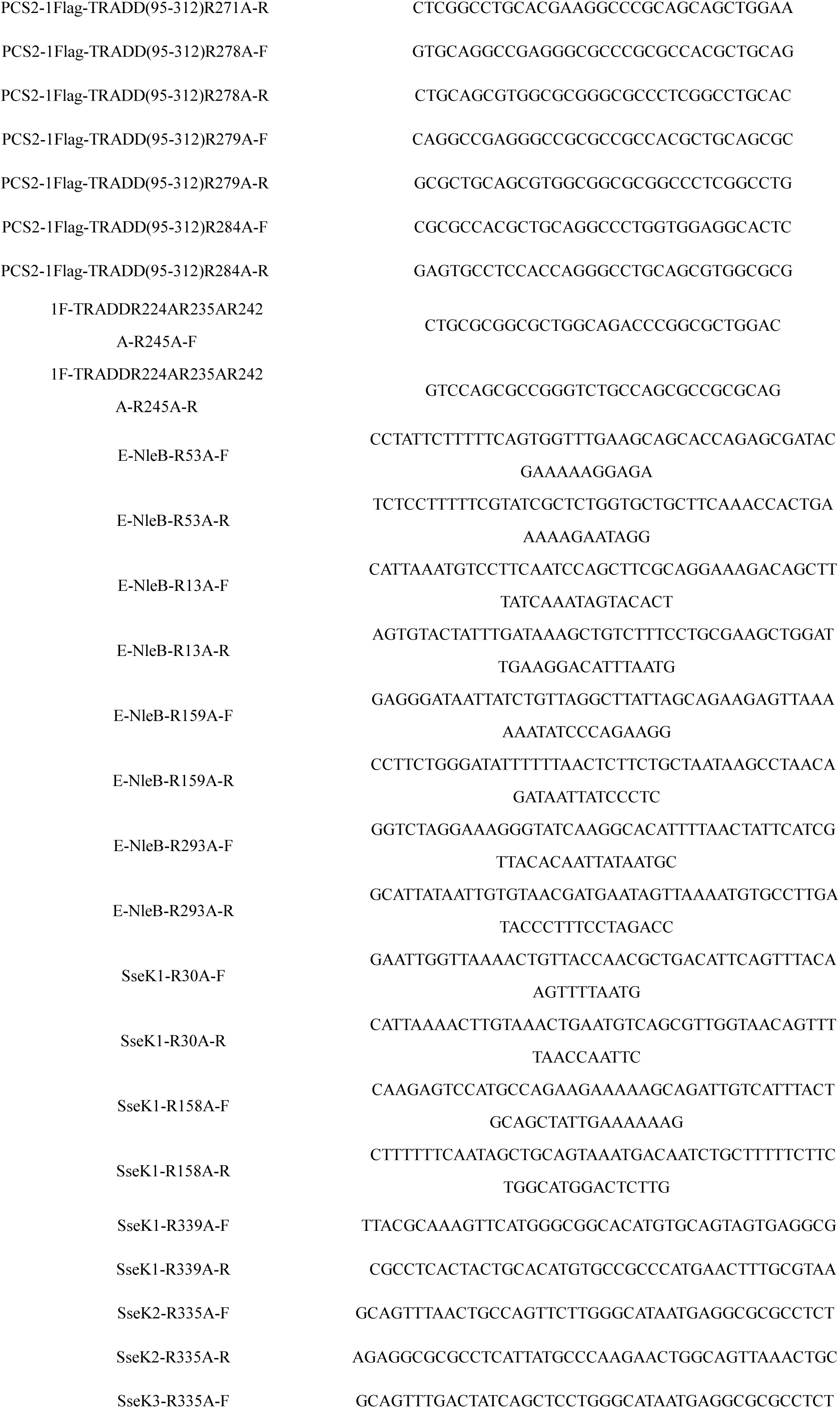
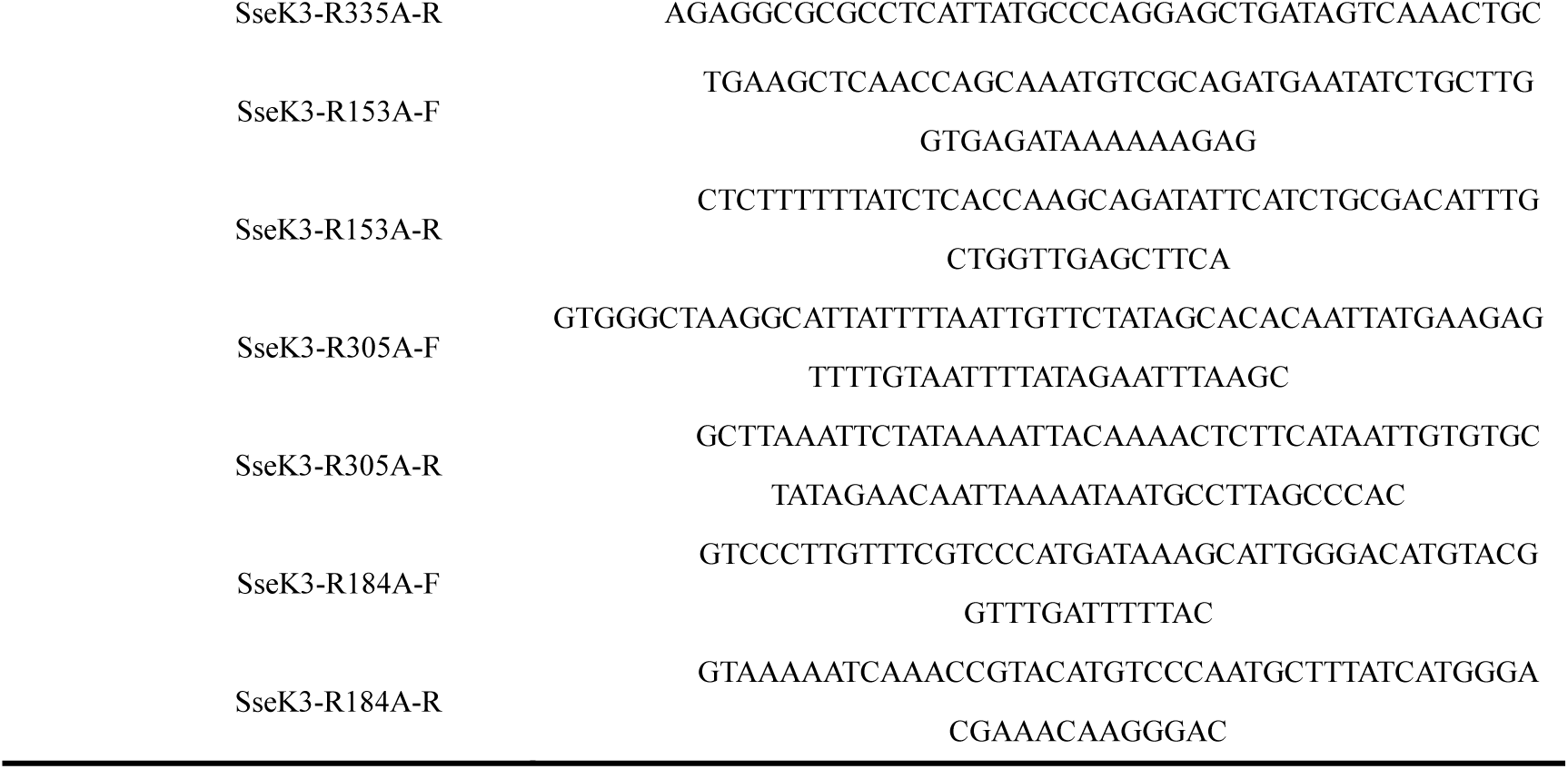
Table Primers used in this study.

